# Trauma Under Psychedelics: How Psychoactive Substances Impact Trauma Processing

**DOI:** 10.1101/2024.03.28.587237

**Authors:** Ophir Netzer, Noa Magal, Yonatan Stern, Tzuk Polinsky, Raz Gross, Roee Admon, Roy Salomon

**Author notes:** These authors contributed equally (Roee Admon, Roy Salomon).

## Abstract

Traumatic events play a causal role in the etiology of stress-related psychopathologies such as depression and posttraumatic stress disorder (PTSD). Recent research highlighted the therapeutic potential of psychoactive substances in alleviating trauma symptoms among chronic stress-related patients. This study is the first to investigate the impact of psychoactive substances consumed during actual trauma exposure. Our cohort includes 772 adult survivors (487 males, Mean±SD Age: 26.96±6.55) of the high-casualty attack that occurred at the Supernova festival in Israel on October 7th, 2023. Survivors completed the study during the peritraumatic period of one to four months following the attack. Primary outcomes include the PTSD Checklist for DSM-5 (PCL-5), the Kessler Psychological Distress Scale (K6), and a self-reported rating of feeling overwhelmed. Secondary outcomes include subjective experiences during the attack and reports on social interactions and sleep quality. All survivors reported being in direct danger of death during the attack. Approximately two-thirds of the sampled survivors were under the influence of psychoactive substances at the time of the attack, including LSD, MDMA, Ketamine, Cannabis, and Alcohol, creating a tragic and unique natural experiment to study the impact of psychoactive compounds on trauma processing. Analysis revealed that participants that were under the influence of MDMA during the attack (n=99) reported feeling less overwhelmed, having more social interactions, improved sleep quality, and reduced psychological distress compared to those not under the influence of any substance during the attack (n=216). In contrast, those consuming Cannabis and/or Alcohol during the attack (n=68) showed higher psychological distress, more PTSD symptoms, and worse sleep quality compared to those not under the influence of any substance during the attack. Together, these novel findings suggest that trauma exposure under the influence of MDMA is associated with reduced psychological distress and higher sociality, possibly mediated through MDMA’s known effects of reducing negative emotions and elevating prosociality, while Cannabis and/or Alcohol consumption produces deleterious effects. Further research is needed to explore the cognitive and physiological mechanisms linking psychoactive substances to trauma recovery and establish the putative protective role of MDMA.

## Introduction

A traumatic event (TE) is defined as exposure to actual or threatened death, serious injury, or sexual violence^1^. Psychophysiological research revealed pronounced heterogeneity in physiological, cognitive, and emotional responses to acute trauma, as well as its impact on mental health. Most individuals show resilience after encountering substantial trauma, yet some develop chronic and debilitating psychopathologies, most commonly depression and posttraumatic stress disorder (PTSD)^2–5^.

Trauma-related psychopathologies are difficult to treat^6–8^. Recent research has focused on how psychoactive substances may assist in the treatment of PTSD and depression^9–12^. Several psychoactive substances, including psilocybin, LSD, ketamine, and 3,4-methylenedioxymethamphetamine (MDMA), have demonstrated efficacy in alleviating mental health problems^13,14^. Among these psychoactive compounds, MDMA-assisted psychotherapy has received substantial attention in the treatment of chronic PTSD^11,15–19^. MDMA induces the release of presynaptic serotonin [5-hydroxytryptamine (5-HT)], dopamine and norepinephrine^20^. It has been suggested that MDMA facilitates the treatment of PTSD by reducing sensations of fear and threat and inducing prosocial feelings and behaviors^11,21^. In contrast, consumption of other substances such as Alcohol have been associated with more severe posttraumatic stress symptoms and poorer clinical outcomes following TEs^22,23^.

On October 7^th^ 2023, a surprise large-scale attack was launched on Israel with devastating effects. A nationwide study of 710 Israeli adults found that PTSD prevalence nearly doubled post-attack, with direct exposure being a key predictor^24^. The Supernova music festival was a target of the attack, with over 360 of the 4000 festival attendees murdered and dozens taken hostage to Gaza. The survivors experienced and witnessed acute, life-threatening traumatic events for an extended duration. Critically, many attendees were under the influence of psychoactive substances during the attack, while other attendees were not under the influence of any substances. This is an unprecedented event in which mass trauma was experienced while in altered states of consciousness. To our knowledge, there is no previous understanding of the immediate or long-term consequences of trauma that is acutely experienced under psychedelics.

Studying the survivors of this tragic natural experiment could provide novel insights into how trauma is experienced under psychedelics and how this may impact trauma processing in the peritraumatic period and determine clinical outcomes. We investigated a large cohort (n=772) of survivors of the Supernova music festival, assessing their acute experiences during the trauma as well as their processing of the traumatic events and clinical status during the peritraumatic period one to four months following the event.

## Methods

### Participants

The study recruited 1239 individuals from an estimated survivor population of approximately 3500, with eligibility criteria requiring self-reported direct exposure to life-threatening danger and completion of the survey within the peritraumatic period. The final cohort consisted of seven hundred seventy-two (n=772) adults (487 males, Mean±SD Age: 26.96±6.55), who survived the attack at the Supernova music festival. These individuals were identified and recruited through a combination of methods, including a collaboration with “Safeheart”, a non-profit organization (NGO) supporting survivors, word-of-mouth among survivors, and via survivors’ support groups on social media platforms. Prior to data collection, informed consent was obtained from all participants. All procedures were performed in compliance with the institutional guidelines of the University of Haifa and have been approved by the Institutional Ethics Committee (approval no. 374/23).

### Data collection

Participant responses were collected from November 2^nd^, 2023, to February 21^st^, 2024, a period extending from 26 to 137 days following the TE. This timeframe was selected to capture the acute and immediate reactions of survivors.

### Survey Design

The survey assessed constructs categorized into three time frames: subjective experiences during the TE, trauma processing in the peritraumatic period, and current clinical outcomes (Fig. 1A). Participants reported demographics (age, sex at birth), level of exposure to traumatic events, and substance use before the TE. Constructs assessed included feelings of control, isolation, substance helpfulness, social support, guilt, sleep quality, and emotional overwhelm, all rated on a 0–100 scale. The Kessler Psychological Distress Scale (K6)^25^ and the PTSD Checklist for DSM-5 (PCL-5)^26^ assessed current levels of mental distress and PTSD symptom severity, respectively. Full survey details are in Supplementary Table S1.

**Figure 1:**
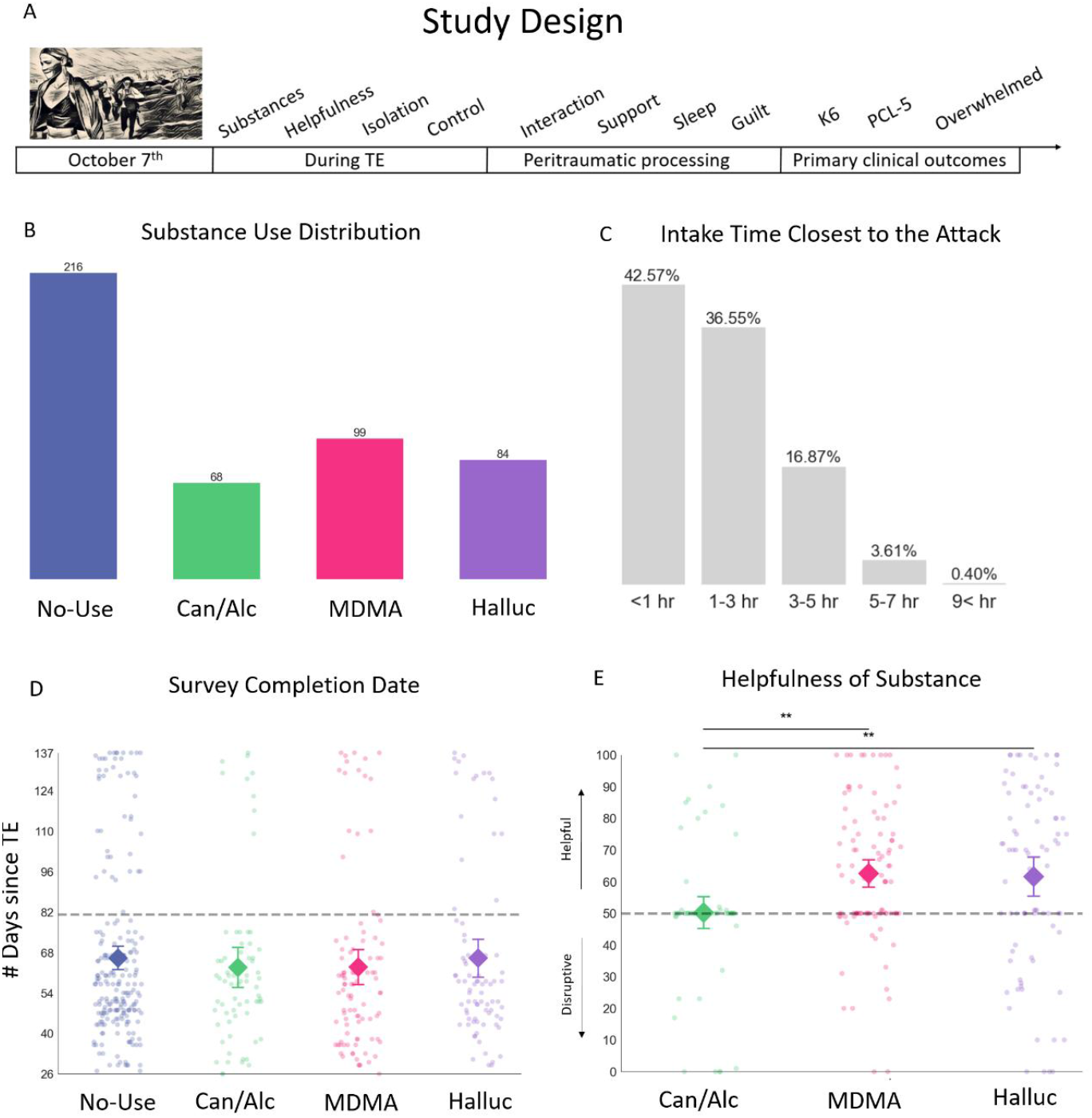
Study design and substance groups distributions. (A) Study design. Schematic representation of the timeline, beginning with the traumatic event on Oct 7^th^, and questions pertaining to subsequent phases; During TE: substance use during the TE, perceived helpfulness, feelings of isolation, and feelings of control; Peritraumatic processing: social interactions, feelings of support, sleep quality and feelings of guilt; Primary clinical outcomes: survivors’ current clinical state including mental distress (K6), PTSD symptoms severity (PCL-5) and feelings of being overwhelmed. All measurements were collected simultaneously but pertain to different epochs of the traumatic event and peritraumatic period. (B) Substances distribution. The majority of participants reported no substance use (n=216), followed by those who were under the influence of MDMA (n=99), Hallucinogens (n=84), and Cannabis and/or Alcohol (n=68). (C) Distribution of last substance intake times relative to the TE. Most participants reported consuming substances within the 3 hours prior to the start of the attack (42.57% within 1 hr, 36.55% within 1-3 hrs). Panels D-E: Individual participants are represented by dots with overlaying mean values (diamonds) and 95% confidence intervals (error bars) for the respective measures in relation to substance groups (*No-Use, Can/Alc, MDMA, Halluc*). (D) Distribution of survey response collection date by substance group. Dots represent the survey completion date for each participant, measured in days from the TE. No differences were found between groups in the time that passed from the TE until survey completion date. (E) Participants’ perceived helpfulness of the substance during the TE. Given no specific hypotheses, an exploratory test comparing all groups against each other was conducted. Dots represent individual helpfulness ratings on a scale between 0 to 100, where 0 indicates the substance was very disruptive and 100 indicates the substance was very helpful. Higher scores represent more perceived helpfulness, with the dashed line indicating the threshold for positive judgments. Notably, individuals who consumed Cannabis and/or Alcohol experienced it as significantly less helpful to them during the TE compared to individuals who consumed MDMA or Hallucinogens (all p<0.05).

### Analytical Approach

Participants were grouped by the substance they reported being under its influence during the attack: *Hallucinogens* (e.g., psilocybin and LSD), *MDMA* (i.e., MDMA and Ecstasy), *Stimulants* (e.g., cocaine and amphetamines), *Can/Alc* (Cannabis and/or Alcohol), *Other* substances (e.g., Ketamine, PCP), and a *No-Use* group (participants that were not under the influence of any substance during the TE). Due to the prevalence of poly-substance use (see Supplementary Table S2 for all combinations) and the focus on isolating the psychopharmacological effects of individual substances, the main analyses included only participants using a single substance without mixtures. Groups with insufficient participant numbers to achieve meaningful statistical power (i.e., n<30) were excluded. Consequently, the primary analyses focused on the following groups of participants: *No-Use* (n=216), *Can/Alc* (n=68), *MDMA* (n=99), and *Halluc* (n=84). The *Can/Alc* group includes survivors who consumed cannabis, alcohol, or both, as no significant differences were observed across main outcome measures. This grouping was informed by prior research^27^, survivor reports, and current data analyses; detailed results are in the Supplementary material. To account for the potential combined effects of substances, analyses of all substance groups, including mixtures, are provided in the Supplementary Results.

### Statistical Analysis

Linear regression analyses evaluated group differences in substance effects on: (A) trauma exposure, (B) peritraumatic processing, (C) primary clinical outcomes. Two models were tested: one with only substance groups and another including age, sex, and time elapsed since the TE. Models with the lowest Bayesian Information Criterion (BIC)^28,29^ and its results are presented in the main text, with both models detailed in the Supplementary Results. Substances effects were categorized as “beneficial” or “detrimental” indicating outcomes better or worse, respectively, compared to the *No-Use* group. All results were based on two-tailed regression analyses against the *No-Use* reference group. Primary outcomes were analyzed using t-tests for group means compared to clinical cutoffs, and partial Chi-square tests for groups proportions differences. Multiple comparisons were corrected using False Discovery Rate (FDR)^30^.

Relative Risk (RR)^31^ quantified the likelihood of favorable outcomes for substance groups compared to the *No-Use* group.

## Results

### Trauma exposure across participants

Among the 772 survivors who completed the survey, the magnitude of trauma exposure was universally high. All participants (100%) reported direct exposure to life-threatening events during the attack, underscoring the severe and immediate threat to their lives. Furthermore, a significant majority (83%) reported witnessing dead or injured individuals during the attack, 80% reported their loved ones were murdered, and 69% reported that loved ones were injured. Critically, all participants met Criterion-A for PTSD as in DSM-5^1^.

### Substance use during trauma exposure

Seventy-two percent (n=556) of survivors reported consuming at least one substance during the festival. MDMA was the most commonly used substance, with 12.8% of participants reported consuming only MDMA and 14.0% reported consuming MDMA with other substances, followed by Hallucinogens, with 10.9% reported consuming only Hallucinogens and 13.9% reported consuming Hallucinogens with other substances, Cannabis and/or Alcohol (*Can/Alc*, 8.8%), Stimulants (*Stim*, 7.5%) and other substances (4.2%) (Fig. 1B). Slightly less than a third of participants (28.0%) reported no substance use (*No-Use* group). Multi-substance use was common, typically with survivors reporting consumption of Cannabis and/or Alcohol in addition to Hallucinogens or MDMA. For a detailed report of all substance combinations see Supplementary Table S2. Most individuals (79.1%) consumed the substance within three hours before the attack (Fig. 1C), presumably to obtain peak effects at sunrise, and reported feeling strongly affected by the substance when the attack began. Analysis also revealed differences in age among the groups (F(3,463)=2.9, p=0.04,*R*^2^=0.02) driven by younger ages in the *MDMA* group (25.7±5.2) compared to the *No-Use* group (28.1±8.1). Furthermore, the analysis revealed a significant difference in sex distribution between groups 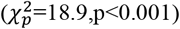, implying that sex may have played a role in the choice of substance. This difference was attributed to a higher proportion of males in the *MDMA* and *Halluc* groups compared to the *No-Use* group (all *p*_*adjusted*_ < 0.01). For the full analysis see the Supplementary Results. No differences between groups were found in the time that passed from the TE until the survey completion date (F(3,463)=0.4, p=0.76, Fig. 1D). Table 1 provides an overview of age, sex and substance use experience. Importantly, possible differences in trauma exposure between groups were assessed using Chi-square test on the proportion of exposure between groups for each variable. Analyses revealed no differences in rates of trauma exposure between groups, across all trauma types (all p-values>0.15). For a detailed account of group-specific exposure rates, see Supplementary Table S5. Chi-square analyses for differences in exposure rates are available in Supplementary Table S6.

**Table 1:**
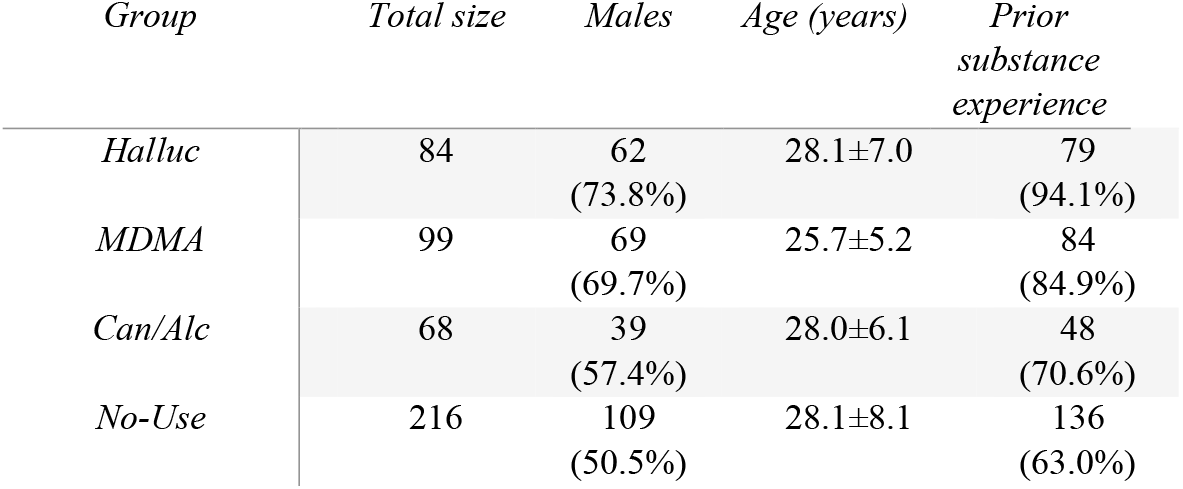
Summary of demographic variables and prior substance experience across substance groups. Variables include sex, age (Mean±SD), and reported prior substance experience. Percentages in parentheses reflect the proportion of participants for each variable relative to the total number in their respective groups. It should be noted that not all participants provided responses for every variable. For detailed information on the distribution of responses refer to Supplementary Table S8.

### Effects of substance during trauma exposure

Surprisingly, many survivors who reported using various substances during the TE described them as generally helpful (Overall Mean±SD:58.9±24.4). A significant difference emerged between groups regarding the perceived influence of the substances during the TE (F(2,247)=6.14, p=0.003, *R*^2^=0.05). Specifically, individuals in the *MDMA* (62.6±21.7,β=12.4,p=0.001) and the *Halluc* (61.5±28.3,β=11.3,p=0.004) groups felt that the substance provided more assistance during the TE compared to the *Can/Alc* group (50.2±20.6; Fig. 1E). Contrary to expectations, no significant differences were found between groups in judgments regarding feelings of control (F(3,463)=0.59,p=0.62) or feelings of isolation (F(3,643)=1.53,p=0.21) during the TE (See Supplementary Fig. S1).

### Effects of substance on peritraumatic processing

Regarding constructs related to the trauma processing following the TE, significant differences emerged between substance groups. The extent of subsequent social interactions differed significantly between groups (F(3,463)=4.5,p=0.004,*R*^2^=0.03; Fig. 2A). The *MDMA* group reported significantly higher levels of social interactions (76.5±26.2) compared to the *No-Use* group (66.5±27.1,β=10.0,p=0.003). The feeling of support from friends and family was overall high (Overall Mean±SD:75.5±24.9) but did not differ significantly between groups (F(3,463)=2.5,p=0.056,*R*^2^=0.016; Fig. S1). Sleep quality was generally poor (Overall Mean±SD:37.4±29.5) and varied significantly between groups (F(3,463)=4.6,p=0.004,*R*^2^=0.03; Fig. 2B). Compared to the *No-Use* group (37.5±30.0), the *MDMA* group reported significantly improved sleep quality (45.4±30.9,β=7.9,p=0.025), while the *Can/Alc* group reported worse sleep (29.4±25.0,β=-8.1,p=0.046). No significant differences were observed in feelings of guilt (F(3,463)=0.97,p=0.41; Fig. 2C), suggesting that substance type during the TE did not influence this aspect of trauma processing.

**Figure 2:**
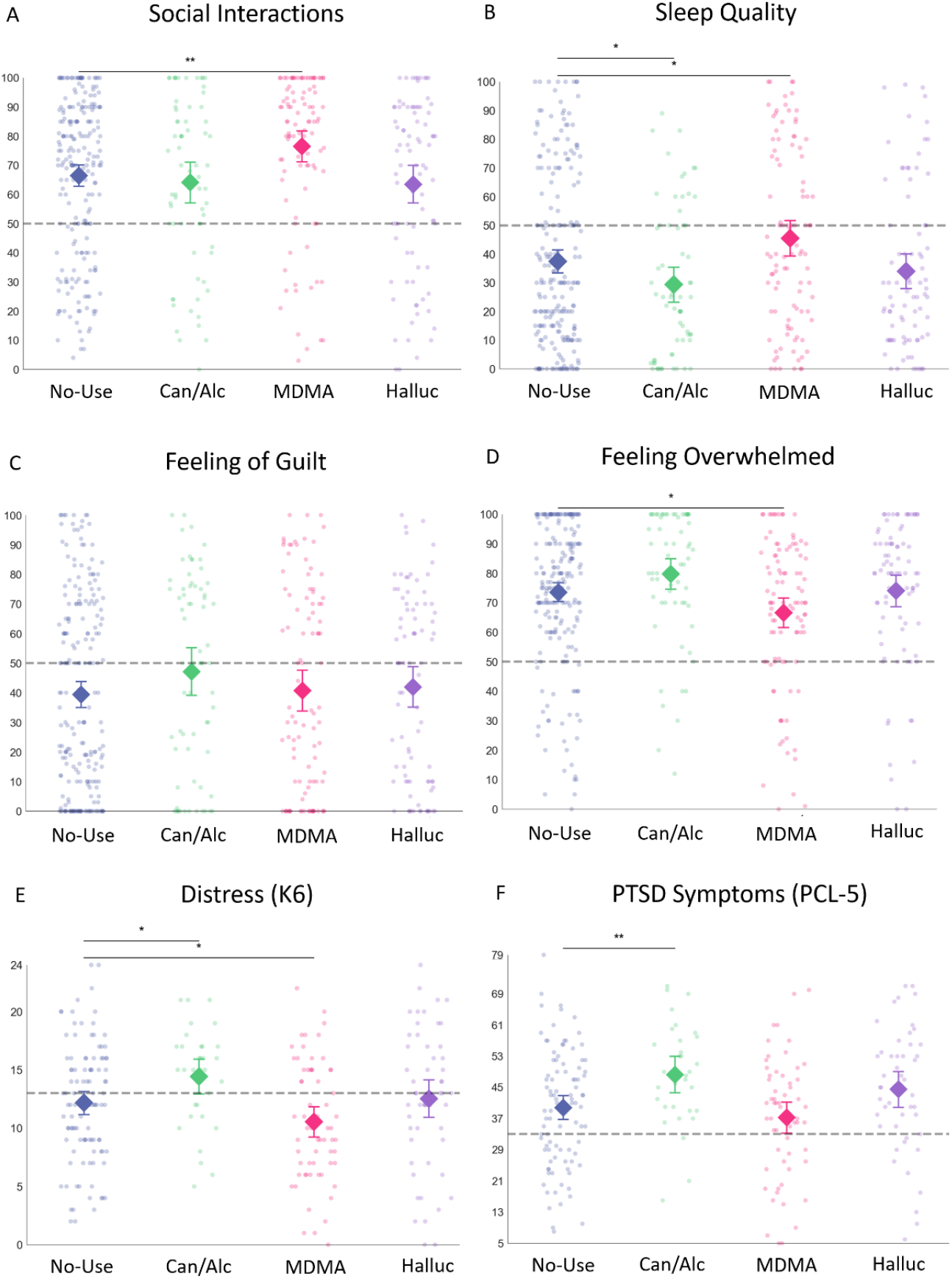
Outcome measures across substance groups. Panels A-F represent dot plots with overlaying mean values (diamonds) and 95% confidence intervals (error bars) for the respective outcome measures in relation to substance groups (*No-Use, Can/Alc, MDMA, Halluc*). Statistical significance is denoted by asterisks (*p < 0.05, **p < 0.01). (A) Social Interactions. Mean levels of perceived social interactions across groups, with the dashed line representing the threshold for positive judgments, illustrating significantly higher levels of social interactions in the *MDMA* group compared to the *No-Use* group. (B) Sleep Quality. Mean levels of sleep quality scores, with the *MDMA* group reporting significantly better sleep quality compared to the *No-Use* group, while the *Can/Alc* group reporting significantly worse sleep quality compared to the *No-Use* group. (C) Feelings of Guilt. Mean levels of reported guilt feelings were analyzed across groups, with no significant differences observed. (D) Feeling Overwhelmed. The reported feeling of being currently overwhelmed, with the *MDMA* group reporting significantly lower levels compared to the *No-Use* group. (E) Distress (K6). The Kessler Psychological Distress Scale (K6) scores, with a dashed line representing the clinical cutoff (=13). The *MDMA* group demonstrated significantly lower distress scores compared to the *No-Use* group, while the *Can/Alc* group reported significantly higher distress compared to the *No-Use* group. (F) PTSD Symptoms (PCL-5). Scores from the PTSD Checklist for DSM-5 (PCL-5), with a dashed line representing the clinical cutoff (=33). The *Can/Alc* group reported significantly higher PCL-5 scores compared to the *No-Use* group, indicating more severe PTSD symptoms.

### Primary Clinical Outcomes

Assessing the current state of participants one to four months following the TE revealed a significant difference between groups in the feeling of being currently overwhelmed (F(3,463)=4.22,p=0.006,*R*^2^=0.03; Fig. 2D). This effect was driven by significantly lower ratings of feeling overwhelmed in the *MDMA* group (66.6±25.2) compared to the *No-Use* group (73.5±23.8,β=-7.0,p=0.017). Mental distress (K6) scores also differed significantly between groups (F(3,250)=4.3, p=0.006,*R*^2^=0.05; Fig. 2E). Here as well, effects were driven by significantly lower K6 scores in the *MDMA* group (10.5±5.1) compared to the *No-Use* group (12.2±5.1; β=-1.6,p=0.049), as well as by significantly higher K6 scores in the *Can/Alc* group (14.4±4.1) compared to the *No-Use* group (β=2.3,p=0.031). Finally, PTSD symptom severity (PCL-5) scores also differed significantly between groups (F(3,229)=4.8,p=0.003,*R*^2^=0.06; Fig. 2F), with significantly higher PCL-5 scores in the *Can/Alc* group (48.3±12.8) compared to the *No-Use* group (39.8±14.9,β=8.5,p=0.006). Notably, mean PCL-5 scores across all groups were high (41.3±15.3), with every group significantly exceeding the clinical cutoff score (*t*_*No*−*Use*_(93) = 4.4, *t*_*Can/Alc*_(31) = 6.6, *t*_*MDMA*_ (58) = 2.1, *t*_*Halluc*_ (47) = 5.1, *p* < 0.02 for all; Clinical cutoff=33). This was not the case with respect to mental distress where the *MDMA* and *No-Use* groups exhibited significantly lower K6 scores than the clinical cutoff, while the *Can/Alc* group was the only group to have significantly higher scores compared to the clinical cutoff (*t*_*No*−*Use*_(103) = −1.7, *t*_*MDMA*_(62) = −3.8, *t*_*Can/Alc*_(32) = 2.0, *p* < 0.05 for all; Clinical cutoff=13).

### Probability analysis for favorable outcomes

Favorable outcomes were defined as the proportion of participants scoring above a positive threshold or below the clinical cutoff. Participants in the *MDMA* group had the highest likelihood of favorable outcomes (Fig. 3), while the *Can/Alc* group had the lowest. Probabilities for the *No-Use* and *Halluc* groups varied across measures. Subsequent RR computations for the primary clinical outcomes comparing substance groups to the *No-Use* group revealed that the *Can/Alc* group had higher risk to exhibit PTSD symptoms above clinical cutoff compared to the *No-Use* group 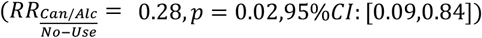. RR analysis of K6 scores and feeling overwhelmed revealed no significant results for any group. For full RR statistics see Supplementary Table S7.

**Figure 3:**
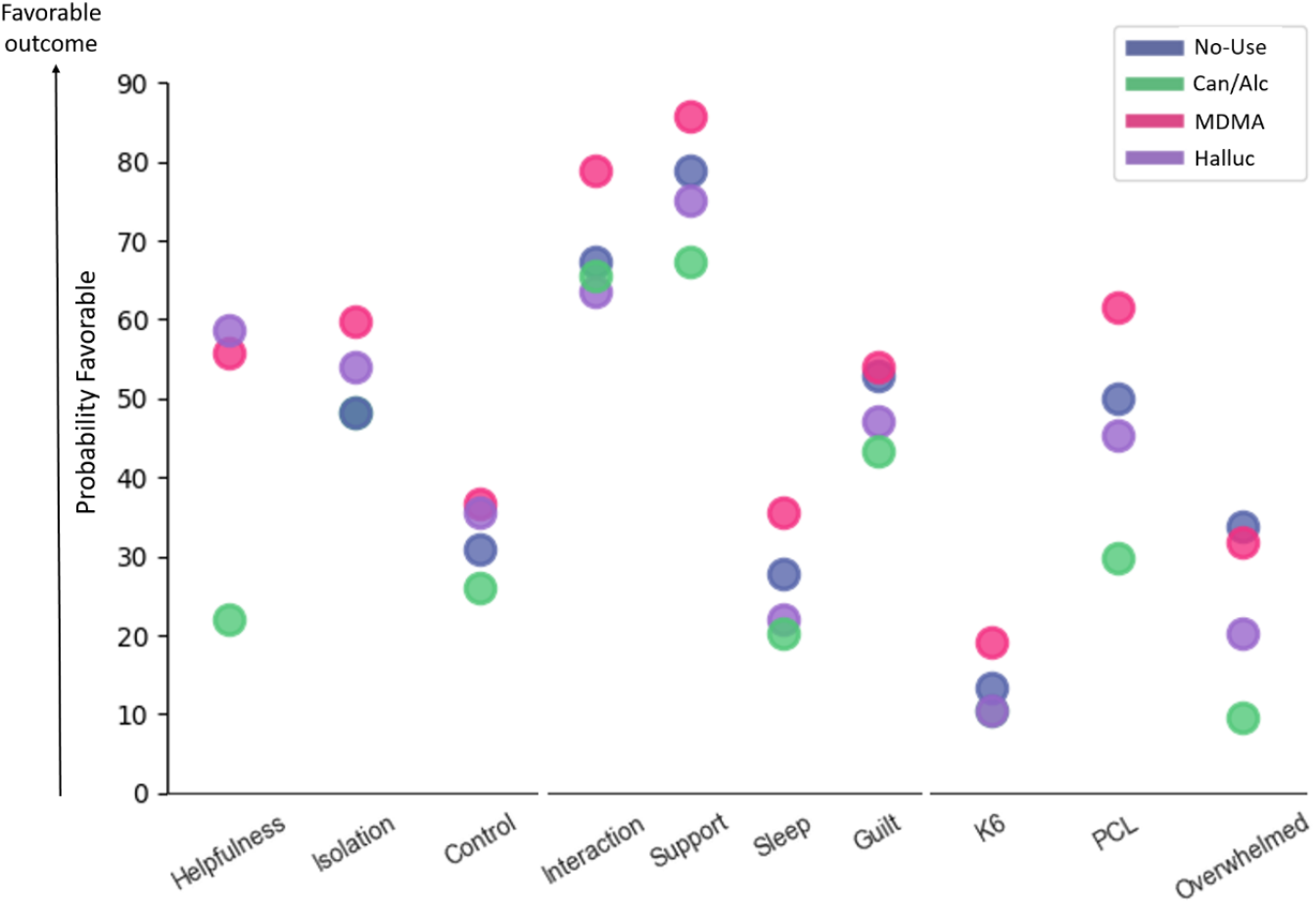
Probability for favorable outcome across substance groups for each trauma related measure. Composite bubble chart depicting the percentage of participants above the cutoff score of each measure in the favorable outcome direction, across different substance groups. Color coding indicates the substance group as per legend. Measures include perceived helpfulness of substance during the TE, feelings of isolation during the TE, feelings of control during the TE, social interaction, support, sleep quality, feelings of guilt during the peritraumatic period, mental distress (K6), PTSD symptom severity (PCL-5), and current feelings of being overwhelmed. The *MDMA* group exhibited more favorable outcomes across almost all measures, while the *Can/Alc* group exhibited less favorable outcomes.

## Discussion

Previous work investigated how psychoactive substances may help alleviate post-traumatic psychopathologies^13,18,32,33^. The Supernova festival attack allowed, for the first time, to investigate the impact of experiencing acute trauma under the influence of psychoactive substances. Our results reveal a novel effect by which survivors of a mass-trauma event who experienced the trauma under the influence of MDMA exhibit enhanced coping across clinical and subjective measures compared to survivors who were not under the influences of substances during the TE. Additionally, survivors who were under the influence of Cannabis and/or Alcohol during the attack showed significantly worse outcomes.

Survivors who consumed MDMA as well as Hallucinogens, reported feeling that the substance assisted them during the TE compared to survivors who consumed Cannabis and/or Alcohol. Anecdotal reports from survivors suggest that the substance may have endowed them with energy to run further and faster during the attack, in line with the known energizing effects of MDMA^34^. These reports also suggest that survivors who were under the influence of MDMA during the trauma may have experienced reduced sensations of fear and threat as the event unfolded.

Additionally, MDMA was associated with improved peritraumatic processing, including increased social interactions compared to survivors who did not consume substance during the TE, aligning with the well-documented prosocial effects of MDMA in both animals and humans^21,35,36^. Interestingly, in the current study, participants in the *MDMA* group exhibited significantly better social coping after the TE, with no differences found on social behavior during the event itself, suggesting that the effect emerged more prominently in the post-trauma period. Thus, the prosocial effects of MDMA may manifest more prominently in the recovery phase.

In terms of clinical outcomes, participants in the *MDMA* group also showed a beneficial effect, with reduced feelings of being overwhelmed and lower mental distress in the months following the trauma, compared to participants who did not consume substances during the TE. This aligns with evidence from MDMA-assisted psychotherapy studies, which highlight reduction of negative affect as key to its therapeutic efficacy^17,37^. Current protocols for MDMA-assisted psychotherapy suggest that re-experiencing traumatic events in a safe setting while experiencing the prosocial and fear-reducing effects of MDMA may enhance the benefits of psychotherapy for PTSD^11,12,36^. The present study extends this idea by demonstrating that even outside a structured therapy setting, MDMA may facilitate adaptive post-trauma social behaviors, potentially supporting psychological recovery and may drive the improved clinical state reported by survivors.

Another construct potentially impacted by MDMA is sleep. Survivors in the *MDMA* group reported better subjective sleep quality over the subsequent one to four months compared to the *No-Use* group. Sleep quality is known to be impaired in the aftermath of trauma and this impairment is considered to not only represent a symptom of trauma exposure, but to play a causal role in the formation of posttraumatic psychopathologies^38–41^. Hence, improved sleep quality in the peritraumatic period may relate to the overall improved clinical state reported by participants in the *MDMA* group.

An additional novel finding of this study is that survivors who were under the influence of Cannabis and/or Alcohol during the TE showed a detrimental effect, with worse outcomes on several scales compared to survivors who did not consume substances. Specifically, survivors who consumed Cannabis and/or Alcohol reported that the substance was less helpful during the TE compared to survivors who consumed each of the other substances. Moreover, they reported significantly worse sleep quality in the peritraumatic period, and poorer clinical outcomes including higher mental distress and worse posttraumatic symptoms compared to participants who did not consume substances during the TE. These findings align with previous research demonstrating the detrimental effects of Alcohol on trauma processing, including increased risk of peritraumatic dissociation, anxiety, depression, and acute stress disorder (ASD) symptoms^42^. Our results extend to previous findings by showing that individuals who consumed Cannabis and/or Alcohol during the TE reported significantly worse sleep quality in addition to more severe clinical symptoms. This may be linked to the established relationship between trauma and sleep disturbances^38–40^. Our results suggest that experiencing trauma under the influence of these substances may exacerbate negative effects during the peritraumatic period.

While considering these novel insights it is important to acknowledge that, as in all natural experiments, we had no control over numerous factors such as participants’ substance choice, dosage, and intake time, as well as potential personality-based biases regarding substance choice. This limitation introduces potential confounding factors. However, this study captures an unprecedented natural experiment, offering unique insights unavailable in controlled studies. The real-world context provides an opportunity to observe trauma processing in mass-trauma survivors who used psychoactive substances during the TE, revealing complex real-life behavior and substance use patterns. Nonetheless, caution is advised when drawing causal inferences from the findings. Additionally, current results pertain to the peritraumatic period, which, while predictive of later posttraumatic psychopathologies, does not capture the full clinical picture, as symptoms have yet to stabilize. Another limitation is merging Cannabis and Alcohol into one group. Although our analyses detected no significant differences between groups, the small sample size prevented us from exploring the distinct effects of each substance individually. Finally, potential biases must be considered. First, a clear survivor bias exists as we cannot collect data from those who tragically did not survive the attack. Second, survivors with more severe symptoms may be underrepresented in our sample. However, the mean PCL-5 score in our sample across groups is well above the clinical cutoff (Mean±SD: 41.29±15.30) suggesting that this cohort shows substantial posttraumatic symptoms. Further research should explore the mechanisms linking psychoactive substances to trauma recovery and establish the putative protective role of MDMA and detrimental effect of Cannabis and Alcohol.

## Conclusions

In summary, this tragic natural experiment provides the first-ever investigation into how psychoactive substances impact trauma processing during acute trauma and the peritraumatic period. Survivors under the influence of MDMA during the TE exhibited better processing and improved clinical outcomes, while those under Cannabis and/or Alcohol exhibited worse processing and more severe clinical states compared to survivors who were not under the influence of substances during the TE. These results are a first report from an ongoing longitudinal study aiming to elucidate the cognitive, physiological, and neural mechanisms underlying the effects of psychoactive substances on trauma processing. We hope our findings will deepen understanding of trauma under psychoactive substances and aid survivors of this harrowing experience.

## Acknowledgments

The authors wish to thank the many people who made this project possible during difficult days in Israel. First, the SafeHeart NGO management and early volunteers (Reut Plonsker, Guy Simon, Karina Dessau, Tal Zagursky, Yair Grynbaum, Nir Tadmor, Igal Tartakovsky, Irit Hacmun, Stephanie Cohen, Shiran Maor, Dr. Demian Halperin & Efrat Atun) who worked tirelessly to ensure treatment for the survivors. We thank all the SafeHeart volunteers, clinicians and the research team. Finally, we wish to extend our deepest gratitude to the amazing survivors of the festival who gave us their trust and made a tremendous effort to help others even in such difficult times.

## Notes

### Competing Interest Statement

The authors have declared no competing interest.

### Summary of Updates

In this revised manuscript, we have made several key updates. First, we have expanded our analyses to include the Hallucinogens group, which encompasses classic psychedelics such as LSD and psilocybin, allowing for a more comprehensive examination of substance-related effects. Second, we have incorporated additional participants identified within the peritraumatic period, increasing the robustness of our findings. Third, to better isolate the psychopharmacological effects of individual substances, our main analyses now focus on participants who reported using a single substance without mixtures. Lastly, to account for potential combined effects of substances, we have included additional analyses incorporating all substance groups, including mixtures, in the supplementary material. These revisions enhance the clarity, specificity, and robustness of our findings while ensuring a more rigorous interpretation of substance-related effects.

