## Supplementary material for "Trauma Under Psychedelics: How Psychoactive Substances Impact Trauma Processing"

#### Methods

##### *Participants*

The study recruited 1239 individuals from an estimated survivor population of approximately 3500, with eligibility criteria requiring self-reported direct exposure to life-threatening danger and completion of the survey within the peritraumatic period. The final cohort consisted of seven hundred seventy-two (n=772) adults (487 males, Mean±SD Age: 26.96±6.55), who survived the attack at the Supernova music festival. These individuals were identified and recruited through a combination of methods, including a collaboration with “Safeheart”, a non-profit organization (NGO) supporting survivors, word-of-mouth among survivors, and via survivors’ support groups on social media platforms. Prior to data collection, informed consent was obtained from all participants. All procedures were performed in compliance with the institutional guidelines of the University of Haifa and have been approved by the Institutional Ethics Committee (approval no. 374/23).

##### *Data collection*

Participants' responses were collected from November 2nd, 2023, to February 21st, 2024, a period extending from 26 to 137 days following the TE. This timeframe was selected to capture the acute and immediate reactions of survivors, thereby providing a snapshot of their initial state. Collecting data in this time window was critical in ensuring the analyses of peritraumatic processes, providing a comprehensive understanding of the survivors' psychological states in the aftermath of the traumatic experience. During this period, participants were contacted via telephone, text message, email, or social media platforms, based on contact information provided by the participants during the initial recruitment stage. The survey was advertised broadly to minimize bias, with the general purpose stated as: "The purpose of this survey is to see how you are doing following the event you experienced". When addressing substance use during the festival, we added within the survey: "We wish to evaluate if and how the substances affected your experience". Each participant completed the survey only once. During this single session, participants were asked to reflect on different time periods since the event and up to date. The survey instrument was hosted on the Qualtrics platform, a secure online system that allows for anonymous and confidential submission of responses.

##### *Survey Design*

Several constructs pertinent to the TE and its subsequent processing were collected. These constructs were categorized into three distinct time frames: constructs related to the subjective acute experiences during the TE itself, constructs relevant to trauma processing in the peritraumatic period following the TE, and the primary clinical outcomes which portrayed the current clinical state of survivors. Participants first provided basic demographic details, including age and sex assigned at birth. They then were queried about their level of exposure to the threat of death, personal injuries sustained, and harm to loved ones during the TE. Next, participants were inquired regarding substance use during the festival preceding the TE, including the type and quantity of substances used and timing of consumption relative to TE onset. Constructs related to the subjective experiences during the TE were also assessed, including feelings of control, feelings of isolation, and the potential helpfulness of substances taken. Measures relevant to peritraumatic processing encompassed social support, social interactions, guilt feelings, and sleep quality since the TE and up to date. Participants rated these constructs on a scale from 0 to 100, with 100 indicating the most extreme manifestation of the construct (e.g., optimal sleep quality or extreme guilt). Finally, subjective ratings of survivors' current state and established psychometric instruments were utilized to measure the primary clinical outcomes. Survivors were asked to rate their current level of feeling emotionally overwhelmed on a scale from 0 to 100, with 100 representing feeling extremely overwhelmed at the present moment. The Kessler Psychological Distress Scale (K6)[1] and the PTSD Checklist for DSM-5 (PCL-5)[2] assessed current levels of mental distress and PTSD symptom severity, respectively. The K6 scale consists of 6 items, with scores interpreted as follows: 0–7 indicates low distress, 8–12 suggests moderate distress, and scores of 13 or higher indicate high distress, often associated with a higher likelihood of mental disorder. The PCL-5 scale includes 20 items assessing PTSD symptoms. A score of 0–32 indicates a lower likelihood of PTSD, while a total score of 33 or higher is commonly used as a cutoff to identify individuals who likely meet the diagnostic criteria for PTSD. Additional data were collected to control for potential confounding variables. Questions targeted the receipt of psychological aid pre- and post-TE, prior trauma experiences, existing psychiatric or neurological diagnoses, chronic medication usage, return to work status post-TE, and prior experience with psychoactive substances. Table S1 outlines all survey questions and clinical assessments that were included in this study. Questions were presented in Hebrew and translated into English to facilitate their inclusion in this manuscript. Responses to all questions were collected simultaneously during a single session, though each question in the table is categorized according to the specific time period it addresses. Furthermore, the table delineates the range of potential answers available for participant in response to each question.

##### *Participants' division to substance groups*

To categorize participants into substance groups, participants were asked to report all substances they consumed during the music festival prior to the terror attack. Participants' responses regarding substance use were assessed and recorded. For analytical clarity, participants were then assigned to distinct groups based on a "winner takes all" methodology. This approach entails classifying each participant into a single group, corresponding to the most potent substance they reported using, irrespective of the number or combination of substances taken. The hierarchy of substance potency, utilized for group allocation, was determined in descending order of presumed impact: Hallucinogens, MDMA, Stimulants, Other substances, Cannabis and/or Alcohol. The rationale behind this ranking system acknowledges the common practice at festivals of poly substance use, where typically one substance predominates in its phenomenological experience and pharmacological effect. Thus, the classification aimed to reflect the most significant substance influence for each participant within the context of the festival environment. To address potential combined effects of substances, we divided participants into smaller groups: those who reported using only the primary substance and those who mixed it with other substances. This division was applied when participant numbers were sufficient to allow meaningful statistical power for analyses, specifically when groups had 30 or more members. As a result, we categorized participants into two groups for hallucinogens: "Hallucinogens", which included individuals who reported using only hallucinogens, and "Hallucinogens polyuse", which comprised those who consumed hallucinogens alongside other substances. Similarly, for MDMA, participants were divided into "MDMA", representing those who exclusively used MDMA, and "MDMA polyuse", for those who used MDMA in combination with other substances. This resulted in the division of the data to the following 8 groups: *Halluc*, *Halluc polyuse, MDMA, MDMA polyuse, Stim, Other, Can/Alc* and *No-Use.* Details of polysubstance use are provided in Table S2.

The supplementary results include two types of analyses. The first, referred to as the 'Main Analysis', focused on participants who reported using a single substance without any mixtures. This analysis corresponds to the primary results detailed in the main text and includes the following groups: *No-Use* (n=216), *Cannabis/Alcohol* (*Can/Alc*, n=68), *MDMA* (n=99), and Hallucinogens (*Halluc*, n=84). The second analysis, termed the 'Comprehensive Analysis', included all substance groups, encompassing participants from both single-substance and polysubstance use categories, with the exception of the "Other" group, which was too small to analyze.

#### *Statistical Analysis*

Linear regression analyses were implemented to assess putative differences between groups with respect to (A) the effects of substances during trauma exposure, (B) the effects of substances on peritraumatic processing, and (C) the effects of substances on the primary outcome measures, reflecting individuals' current clinical state. For each variable, a model that included only the substance groups was initially implemented. Then, a second model was computed accounting for potential confounding factors such as demographics (e.g., age and sex). The two models were compared using Bayesian Information Criterion (BIC[3,4]). Specifically, a BIC difference (ΔBIC) of 0–2 suggests weak evidence, 2–6 indicates positive evidence, 6–10 implies strong evidence, and a difference greater than 10 provides very strong evidence for the model’s improved explanatory power[4]. The supplementary results section depicts the results of both the first model and the second model with additional variables.

An exploratory correlation analysis was conducted to elucidate the relationships between the study constructs. The analysis was conducted both across all main analysis groups combined (*Halluc*, *MDMA*, *Can/Alc*, *No-Use*) and within individual substance groups to explore construct relationships by group. Results are presented in supplementary Fig. S4. Statistically significant correlations ($p<0.05$, unadjusted for multiple comparisons) are marked with asterisks.

**Table S1.** Description of all questions in the online survey included in the current study.

| *Question* | *Section* | *Possible response* |
|---|---|---|
| *Exposure to trauma* | During the TE | Yes / No response for each of the following options: lost consciousness; was in direct danger of death; sustained a major physical injury; sustained a minor physical injury; people that were close got hurt; people that were close were killed; you saw other people injured or dead around you. |
| *Have you taken any substances during the festival* | During the TE | Yes / No / prefer not to say |
| *Which substances have you taken during the festival* | During the TE | Choice from a list of substances with an option of adding a text response. |
| *How long before the start of the attack did you consume the substances* | During the TE | Multiple selection from a list including time ranges: <1h, 1h to 3h, 3h to 5h, 5h to 7h, 7h to 9h, 9> |
| *How strong was the substance's influence before the start of the attack* | During the TE | Continues rating on a scale from 0-100. 0- did not feel the substance influence, 100- felt the substance influence very strongly. |
| *What was the substances' influence during the attack* | During the TE | Continues rating on a scale from 0-100. 0- the substance was very disruptive, 100- the substance was very helpful. |
| *How much control do you feel you had during the attack* | During the TE | Continues rating on a scale from 0-100. 0- did not feel in control at all, 100- felt I was very much in control. |
| *How alone did you feel during the attack* | During the TE | Continues rating on a scale from 0-100. 0- did not feel alone at all, 100- felt I was very alone. |
| *How supported do you feel from your close environment (family, friends)* | Peritraumatic processing | Continues rating on a scale from 0-100. 0- not supported at all, 100- very supported. |
| *How often do you meet people (social interactions)* | Peritraumatic processing | Continues rating on a scale from 0-100. 0- not interacting at all, 100- interacting a lot. |
| *How well do you sleep* | Peritraumatic processing | Continues rating on a scale from 0-100. 0- sleep quality is very poor, 100- sleep quality is very good. |
| *Do you have feelings of guilt* | Peritraumatic processing | Continues rating on a scale from 0-100. 0- not having feelings of guilt at all, 100- feeling a lot of guilt. |

| | | |
|---|---|---|
| *How overwhelmed do you feel right now* | Primary clinical outcome | Continues rating on a scale from 0-100. 0- Not feeling overwhelmed at all, 100- feeling extremely overwhelmed. |
| *K6 questionnaire* | Primary clinical outcome | The K6 scale consists of 6 items, with scores interpreted as follows: 0–7 indicates low distress, 8–12 suggests moderate distress, and scores of 13 or higher indicate high distress, often associated with a higher likelihood of mental disorder. |
| *PCL-5 questionnaire* | Primary clinical outcome | The PCL-5 scale includes 20 items assessing PTSD symptoms. A score of 0–32 indicates a lower likelihood of PTSD, while a total score of 33 or higher is commonly used as a cutoff to identify individuals who likely meet the diagnostic criteria for PTSD. |
| *Have you received psychological aid since the TE* | Additional variables of interest | Yes / No |
| *Have you taken substances in the past* | Additional variables of interest | Yes / No / prefer not to say |
| *Which substances have you taken in the past and how often* | Additional variables of interest | Choice from a list of substances with an option of adding a text response. Next to each selection we asked to specify the estimated frequency. |
| *Do you take chronic medications* | Additional variables of interest | Yes / No |
| *Have you ever been diagnosed with a neurological or psychiatric condition* | Additional variables of interest | Yes / No |
| *Have you ever been to psychological therapy before October 7$^{th}$* | Additional variables of interest | Yes / No |
| *Have you experienced other traumatic events in the past other than the event on October 7$^{th}$* | Additional variables of interest | Yes / No |

### Results

#### *Combinations of substances (polysubstance use)*

Table S2 displays all substance combinations within the study cohort, along with the number of individuals who consumed each combination and the corresponding percentage of the total cohort.

**Table S2.** Substance combinations and consumption rates in the study cohort.

| *Substances* | *Count* | *Percentage* |
|---|---|---|
| *Cannabis* | 16 | 2.07 |
| *MDMA* | 77 | 9.97 |
| *Alcohol,LSD* | 14 | 1.81 |
| *Alcohol,Cocaine* | 6 | 0.78 |
| *Alcohol,MDMA,Cannabis* | 14 | 1.81 |
| *Alcohol,LSD,MDMA,Cannabis* | 5 | 0.65 |
| *Alcohol,Ketamine,MDMA* | 3 | 0.39 |
| *LSD* | 76 | 9.84 |
| *Ecstasy,Ketamine,LSD,2CB* | 1 | 0.13 |
| *Alcohol,LSD,Cannabis* | 14 | 1.81 |
| *MDMA,Cannabis* | 14 | 1.81 |
| *Alcohol,Cannabis* | 31 | 4.02 |
| *Alcohol,catha,Cannabis* | 3 | 0.39 |
| *Alcohol,Ketamine,LSD* | 4 | 0.52 |
| *Alcohol,MDMA* | 13 | 1.68 |
| *Alcohol* | 21 | 2.72 |
| *Alcohol,mescaline,Cannabis* | 1 | 0.13 |
| *MDMA,MMC* | 4 | 0.52 |
| *Ecstasy* | 15 | 1.94 |
| *Cocaine* | 7 | 0.91 |
| *speed* | 1 | 0.13 |
| *Alcohol,Ecstasy,MDMA,Cannabis* | 5 | 0.65 |
| *Ecstasy,MDMA* | 7 | 0.91 |
| *Alcohol,Cocaine,Ketamine,MDMA,Cannabis* | 1 | 0.13 |
| *Ecstasy,Ketamine* | 5 | 0.65 |
| *Alcohol,2CB* | 1 | 0.13 |
| *Dosa,MDMA* | 2 | 0.26 |
| *Ketamine* | 11 | 1.42 |
| *Alcohol,Ecstasy,Ketamine,2CB,Cannabis* | 1 | 0.13 |

| | | |
|---|---|---|
| *Alcohol,Dosa,Ecstasy,Ketamine,MMC,Cannabis* | 1 | 0.13 |
| *Alcohol,Cocaine,LSD* | 1 | 0.13 |
| *Alcohol,Ecstasy,LSD,Cannabis* | 1 | 0.13 |
| *LSD,Cannabis* | 13 | 1.68 |
| *MMC,speed* | 1 | 0.13 |
| *Dosa,MMC* | 1 | 0.13 |
| *Ketamine,MDMA* | 7 | 0.91 |
| *Dosa,Ketamine* | 2 | 0.26 |
| *Alcohol,Ecstasy,Ketamine,speed,Cannabis* | 1 | 0.13 |
| *Dosa* | 8 | 1.04 |
| *Alcohol,Ketamine* | 2 | 0.26 |
| *Alcohol,Ketamine,Cannabis* | 3 | 0.39 |
| *Ketamine,LSD* | 8 | 1.04 |
| *LSD,MDMA* | 6 | 0.78 |
| *Alcohol,Ecstasy,Cannabis* | 1 | 0.13 |
| *Alcohol,LSD,MMC,Cannabis* | 1 | 0.13 |
| *Ecstasy,Ketamine,MDMA,Cannabis* | 2 | 0.26 |
| *Alcohol,Ketamine,MDMA,Cannabis* | 1 | 0.13 |
| *Alcohol,Cocaine,Ketamine,MDMA* | 1 | 0.13 |
| *LSD,Psilocybin* | 3 | 0.39 |
| *Alcohol,Dosa,MDMA* | 3 | 0.39 |
| *Other* | 13 | 1.68 |
| *Ketamine,LSD,Cannabis* | 4 | 0.52 |
| *Psilocybin* | 5 | 0.65 |
| *Ecstasy,Ketamine,Psilocybin,Cannabis* | 1 | 0.13 |
| *Alcohol,Ketamine,LSD,Cannabis* | 5 | 0.65 |
| *Ketamine,LSD,MDMA* | 3 | 0.39 |
| *Alcohol,LSD,Psilocybin* | 1 | 0.13 |
| *Ecstasy,Ketamine,MDMA* | 2 | 0.26 |
| *Alcohol,Cocaine,Ketamine,MMC,Cannabis* | 1 | 0.13 |
| *Alcohol,catha,MDMA,Cannabis* | 1 | 0.13 |
| *Dosa,MDMA,Cannabis* | 1 | 0.13 |
| *Alcohol,LSD,Other,Cannabis* | 1 | 0.13 |
| *MMC* | 7 | 0.91 |
| *Alcohol,Ecstasy,Ketamine* | 2 | 0.26 |
| *Alcohol,Cocaine,MDMA* | 3 | 0.39 |
| *2CB* | 1 | 0.13 |
| *Alcohol,Cocaine,Ketamine,Cannabis* | 1 | 0.13 |
| *Ketamine,MDMA,Cannabis* | 1 | 0.13 |
| *Ecstasy,LSD,Cannabis* | 1 | 0.13 |

| | | |
|---|---|---|
| *Alcohol,Ketamine,MMC* | 1 | 0.13 |
| *Alcohol,Cocaine,Dosa,Ketamine,MDMA,Cannabis* | 1 | 0.13 |
| *MMC,Cannabis* | 3 | 0.39 |
| *Ketamine,MDMA,MMC,Cannabis* | 1 | 0.13 |
| *Cocaine,Ketamine* | 2 | 0.26 |
| *LSD,Psilocybin,Cannabis* | 1 | 0.13 |
| *Dosa,Ketamine,Cannabis* | 1 | 0.13 |
| *Alcohol,Dosa,Ecstasy,Cannabis* | 2 | 0.26 |
| *Alcohol,Psilocybin* | 1 | 0.13 |
| *Dosa,Ketamine,LSD* | 1 | 0.13 |
| *Alcohol,Dosa,Ketamine,MMC* | 1 | 0.13 |
| *Ketamine,MMC,2CB,Cannabis* | 1 | 0.13 |
| *Dosa,LSD,Cannabis* | 2 | 0.26 |
| *Alcohol,Cocaine,Ecstasy* | 1 | 0.13 |
| *Alcohol,Dosa,Ecstasy,Ketamine,Cannabis* | 1 | 0.13 |
| *Alcohol,Ecstasy,Ketamine,MDMA* | 1 | 0.13 |
| *Ketamine,Psilocybin* | 1 | 0.13 |
| *Ketamine,MMC,Cannabis* | 1 | 0.13 |
| *Alcohol,Ecstasy,MMC* | 2 | 0.26 |
| *Alcohol,Cocaine,Cannabis* | 1 | 0.13 |
| *Cocaine,LSD,MDMA* | 1 | 0.13 |
| *Alcohol,Cocaine,Dosa,Ecstasy,Ketamine,MDMA,Cannabis* | 1 | 0.13 |
| *Dosa,Ketamine,LSD,MDMA* | 1 | 0.13 |
| *Dosa,Ecstasy* | 2 | 0.26 |
| *PCP* | 1 | 0.13 |
| *Alcohol,Dosa,LSD* | 1 | 0.13 |
| *Cocaine,Ecstasy,Ketamine,Cannabis* | 1 | 0.13 |
| *Ketamine,MDMA,Psilocybin* | 1 | 0.13 |
| *Cocaine,Cannabis* | 1 | 0.13 |
| *Cocaine,Dosa,Ketamine,MMC* | 1 | 0.13 |
| *Dosa,LSD* | 1 | 0.13 |
| *Alcohol,Cocaine,Ketamine,MDMA,Psilocybin,Cannabis* | 1 | 0.13 |
| *Alcohol,Ecstasy,MMC,Psilocybin* | 1 | 0.13 |
| *Cocaine,Dosa,MDMA,Cannabis* | 1 | 0.13 |
| *Alcohol,Cocaine,Ketamine,MDMA,MMC,Cannabis* | 1 | 0.13 |
| *Cocaine,Dosa* | 2 | 0.26 |
| *Dosa,Ecstasy,MMC* | 1 | 0.13 |
| *Alcohol,Cocaine,DMT,Ketamine,LSD* | 1 | 0.13 |
| *LSD,2CB* | 1 | 0.13 |
| *Alcohol,Cocaine,speed* | 1 | 0.13 |

| | | |
|---|---|---|
| *Alcohol,MMC,Cannabis* | 2 | 0.26 |
| *Ketamine,LSD,2CB* | 1 | 0.13 |
| *LSD,MDMA,Cannabis* | 1 | 0.13 |
| *Ketamine,mescaline* | 1 | 0.13 |
| *Alcohol,Cocaine,MDMA,Cannabis* | 1 | 0.13 |
| *Alcohol,MMC* | 1 | 0.13 |
| *Alcohol,MDMA,MMC* | 1 | 0.13 |
| *Dosa,Ecstasy,LSD,MDMA* | 1 | 0.13 |
| *Alcohol,LSD,MMC* | 1 | 0.13 |
| *Alcohol,Cocaine,MDMA,MMC* | 1 | 0.13 |
| *Ketamine,MMC* | 1 | 0.13 |
| *Alcohol,Cocaine,Psilocybin,Cannabis* | 1 | 0.13 |
| *Ketamine,LSD,MDMA,Cannabis* | 1 | 0.13 |
| *No-Use* | 216 | 27.98 |

##### *Demographics Distribution Across Groups*

Table S3 provides an overview of group size, age and sex for all groups. Within the cohort of this study, males constituted 63.1%, totaling 487 individuals. Disaggregating by subgroup, significant differences were found in sex distribution between all substance groups ($\chi_p^2$=41.05,p<0.0001), implying that sex may have played a role in the choice of substance. This difference was attributed to a higher proportion of males in the following groups: *Halluc*, compared to the *Stim* and *No-Use* groups; *Halluc polyuse*, compared to the *Can/Alc*, *Stim*, and *No-Use* groups; *MDMA*, compared to the *Stim* and *No-Use* groups; and *MDMA polyuse*, compared to the *No-Use* group (all p-values<0.04). This distribution indicates a varied representation of male participants across the different substance groups in the study. Table S4 presents the results of the chi-square test and post hoc corrections. Analysis also revealed differences in age among the groups (F(6,733)=4.05, p=0.0005,$R^2$=0.03) driven by younger ages in the *MDMA* group (25.7±5.2) compared to the *Halluc* (28.1±7.0), *Can/Alc* (28.0±6.1) and the *No-Use* (28.1±8.1) groups, and from younger ages in the *MDMA polyuse* (25.0±5.1) compared to the *No-Use* group, indicating potential variations in substance preference across different age demographics. No differences between groups were found in the time that passed from the TE until the survey completion date (F(6,733)=1.1, p=0.38).

**Table S3**: Summary of demographic variables across substance groups.

| *Group* | *Total Size* | *Males* | *Age (years) Mean +- SD* |
|---|---|---|---|
| *Halluc* | 107 | 62 (73.8%) | 28.1±7.0 |
| *Halluc polyuse* | 84 | 86 (80.4%) | 27.4±6.0 |
| *MDMA* | 108 | 69 (69.7%) | 25.7±5.2 |
| *MDMA polyuse* | 99 | 71 (65.7%) | 25.0±5.1 |
| *Stim* | 58 | 28 (48.3%) | 26.3±4.8 |
| *Can/Alc* | 68 | 39 (57.4%) | 28.0±6.1 |
| *No-Use* | 216 | 109 (50.5%) | 28.1±8.1 |

**Table S4**. Chi-square test results for differences in sex distribution proportions between groups, including original and FDR-adjusted p-values.

| | *Original Chi p-value* | *Adjusted Chi p values:* |
|---|---|---|
| *Can/Alc vs MDMA* | 0.140 | 0.227 |
| *Can/Alc vs Halluc polyuse* | 0.002 | 0.009 |
| *Can/Alc vs Stim* | 0.402 | 0.469 |
| *Can/Alc vs MDMA polyuse* | 0.337 | 0.469 |
| *Can/Alc vs No-Use* | 0.394 | 0.469 |
| *Can/Alc vs Halluc* | 0.050 | 0.095 |
| *MDMA vs Halluc polyuse* | 0.107 | 0.187 |
| *MDMA vs Stim* | 0.013 | 0.034 |
| *MDMA vs MDMA polyuse* | 0.646 | 0.685 |

| | | |
|---|---|---|
| *MDMA vs No-Use* | 0.002 | 0.009 |
| *MDMA vs Halluc* | 0.653 | 0.685 |
| *Halluc polyuse vs Stim* | 0.000 | 0.000 |
| *Halluc polyuse vs MDMA polyuse* | 0.024 | 0.055 |
| *Halluc polyuse vs No-Use* | 0.000 | 0.000 |
| *Halluc polyuse vs Halluc* | 0.366 | 0.469 |
| *Stim vs MDMA polyuse* | 0.043 | 0.091 |
| *Stim vs No-Use* | 0.882 | 0.882 |
| *Stim vs Halluc* | 0.003 | 0.012 |
| *MDMA polyuse vs No-Use* | 0.013 | 0.034 |
| *MDMA polyuse vs Halluc* | 0.296 | 0.444 |
| *No-Use vs Halluc* | 0.000 | 0.003 |

#### *Trauma exposure*

The survivors in this study were all exposed to high levels of trauma. Specifically, all survivors reported their life were in danger (100%), 47 reported they lost consciousness during the attack (6.09%), 39 reported sustaining severe physical injury (5.05%), 232 reported sustaining light physical injury (30.05%), 534 reported their loved ones were injured (69.17%), 615 reported their loved were killed (79.66%), 640 reported seeing injured or killed individuals during the attack (82.90%). Exposure rates per group are specified in the Table S5 below. To examine possible differences in exposure rates across groups we performed a chi-square test. No differences were found in exposure rates distributions between groups. Table S6 presents these results.

**Table S5.** Exposure rates by group.

| *Group* | *Life in danger* | *Lost consciousness* | *Severe physical injury* | *Light physical injury* | *People close were injured* | *People close were killed* | *Saw injured or killed individuals* |
|---|---|---|---|---|---|---|---|
| *Halluc* | 84 (100%) | 2 (2.38%) | 5 (5.95%) | 23 (27.38%) | 65 (77.38%) | 71 (84.52%) | 63 (75.00%) |
| *Halluc polyuse* | 107 (100%) | 4 (3.74%) | 3 (2.80%) | 33 (30.84%) | 76 (71.03%) | 87 (81.31%) | 93 (86.92%) |
| *MDMA* | 99 (100%) | 6 (6.06%) | 8 (8.08%) | 30 (30.30%) | 63 (63.64%) | 71 (71.72%) | 80 (80.81%) |
| *MDMA polyuse* | 108 (100%) | 8 (7.41%) | 5 (4.63%) | 30 (27.78%) | 74 (68.52%) | 86 (79.63%) | 98 (90.74%) |
| *Stim* | 58 (100%) | 3 (5.17%) | 1 (1.72%) | 15 (25.86%) | 36 (62.07%) | 45 (77.59%) | 45 (77.59%) |
| *Can/Alc* | 68 (100%) | 6 (8.82%) | 4 (5.88%) | 23 (33.82%) | 50 (73.53%) | 55 (80.88%) | 54 (79.41%) |
| *No-Use* | 216 (100%) | 12 (5.56%) | 8 (3.70%) | 62 (28.70%) | 149 (68.98%) | 174 (80.56%) | 181 (83.80%) |

**Table S6.** Chi-square test results for exposure rates across groups.

| *Exposure* | *X-squared* | *DoF* | *p-value* |
|---|---|---|---|
| *Lost consciousness** | 4.45 | 6 | 0.62 |
| *Severe physical injury** | 5.62 | 6 | 0.47 |
| *Light physical injury* | 1.48 | 6 | 0.96 |
| *People close were injured* | 6.26 | 6 | 0.40 |
| *People close were killed* | 5.57 | 6 | 0.47 |
| *Saw injured or killed individuals* | 11.80 | 6 | 0.07 |

* This test compares proportions between groups; however, one or more groups have fewer than 5 individuals (n < 5), which may compromise the accuracy of the results. In the main analyses, only tests with more than 5 individuals per group were reported and considered reliable. However, the full results are provided here for completeness.

*Substance intake times closest to the TE*

Most individuals consumed the substance within three hours prior to the attack, presumably to obtain peak effects of the substances during sunrise. Accordingly, individuals across substance groups reported being strongly under the influence of the different substances when the attack began at 6:29 AM. Statistical analysis using a chi-square test revealed a significant difference in substance intake time between groups ($\chi_p^2$=6.7,p=0.04), driven by a higher proportion of survivors in the *Can/Alc* group reported taking the substance closer in time to the beginning of the attack compared to *MDMA* and *Halluc* groups (all p_adjusted<0.02), in line with the shorter duration of effects typically associated with Cannabis and Alcohol.

*Full statistics for all constructs*

For each measure evaluated in the main analysis, we first provide a descriptive statistics table with the mean, standard deviation (SD), sample size (N), and confidence intervals (CI lower and upper) for each group and the overall measure.

<u>*Feelings of Control*</u>

| *Substances group* | *Mean* | *SD* | *N* | *Lower CI* | *Upper CI* |
|---|---|---|---|---|---|
| *Halluc* | 35.6 | 34.8 | 84 | 28.0 | 43.2 |
| *Halluc polyuse* | 36.3 | 34.4 | 107 | 29.7 | 42.9 |
| *MDMA* | 36.8 | 36.6 | 99 | 29.5 | 44.2 |
| *MDMA polyuse* | 38.0 | 38.0 | 108 | 30.7 | 45.3 |
| *Stim* | 30.9 | 28.2 | 58 | 23.4 | 38.4 |
| *Can/Alc* | 29.9 | 33.4 | 68 | 21.8 | 38.0 |
| *No-Use* | 35.5 | 35.1 | 216 | 30.8 | 40.3 |
| *All* | 35.3 | 35.1 | 740 | 32.8 | 37.8 |

<u>*Feelings of Isolation*</u>

| *Substances group* | *Mean* | *SD* | *N* | *Lower CI* | *Upper CI* |
|---|---|---|---|---|---|
| *Halluc* | 46.1 | 35.6 | 84 | 38.3 | 53.8 |
| *Halluc polyuse* | 48.0 | 38.5 | 107 | 40.6 | 55.4 |
| *MDMA* | 38.4 | 36.1 | 99 | 31.2 | 45.7 |
| *MDMA polyuse* | 48.3 | 38.7 | 108 | 40.9 | 55.8 |
| *Stim* | 49.3 | 33.2 | 58 | 40.5 | 58.1 |
| *Can/Alc* | 48.0 | 37.5 | 68 | 38.9 | 57.2 |
| *No-Use* | 47.3 | 36.1 | 216 | 42.4 | 52.1 |
| *All* | 46.4 | 36.8 | 740 | 43.8 | 49.1 |

<u>*Helpfulness*</u>

| *Substances group* | *Mean* | *SD* | *N* | *Lower CI* | *Upper CI* |
|---|---|---|---|---|---|
| *Halluc* | 61.5 | 28.3 | 84 | 55.4 | 67.7 |
| *Halluc polyuse* | 62.5 | 27.5 | 106 | 57.2 | 67.8 |
| *MDMA* | 62.6 | 21.7 | 98 | 58.2 | 67.0 |
| *MDMA polyuse* | 60.7 | 22.3 | 108 | 56.4 | 65.0 |
| *Stim* | 61.0 | 21.1 | 58 | 55.4 | 66.6 |

| | | | | | |
|---|---|---|---|---|---|
| *Can/Alc* | 50.2 | 20.6 | 68 | 45.2 | 55.2 |
| *All* | 60.2 | 24.4 | 522 | 58.1 | 62.3 |

*Social Interactions*

| *Substances group* | *Mean* | *SD* | *N* | *Lower CI* | *Upper CI* |
|---|---|---|---|---|---|
| *Halluc* | 63.5 | 29.6 | 84 | 57.1 | 70.0 |
| *Halluc polyuse* | 65.5 | 28.4 | 107 | 60.0 | 71.0 |
| *MDMA* | 76.5 | 26.2 | 99 | 71.3 | 81.8 |
| *MDMA polyuse* | 74.2 | 25.0 | 108 | 69.4 | 78.9 |
| *Stim* | 68.5 | 26.8 | 58 | 61.4 | 75.6 |
| *Can/Alc* | 64.1 | 28.7 | 68 | 57.1 | 71.1 |
| *No-Use* | 66.5 | 27.1 | 216 | 62.9 | 70.1 |
| *All* | 68.4 | 27.7 | 740 | 66.4 | 70.4 |

*Feeling of Support*

| *Substances group* | *Mean* | *SD* | *N* | *Lower CI* | *Upper CI* |
|---|---|---|---|---|---|
| *Halluc* | 72.6 | 26.6 | 84 | 66.8 | 78.4 |
| *Halluc polyuse* | 73.0 | 27.7 | 107 | 67.6 | 78.3 |
| *MDMA* | 80.0 | 22.9 | 99 | 75.4 | 84.6 |
| *MDMA polyuse* | 85.1 | 19.4 | 108 | 81.4 | 88.8 |
| *Stim* | 77.7 | 22.9 | 58 | 71.7 | 83.8 |
| *Can/Alc* | 70.3 | 25.0 | 68 | 64.2 | 76.4 |
| *No-Use* | 76.3 | 24.7 | 216 | 73.0 | 79.6 |
| *All* | 76.7 | 24.7 | 740 | 74.9 | 78.5 |

*Sleep Quality*

| *Substances group* | *Mean* | *SD* | *N* | *Lower CI* | *Upper CI* |
|---|---|---|---|---|---|
| *Halluc* | 34.0 | 27.6 | 84 | 28.0 | 40.0 |
| *Halluc polyuse* | 32.7 | 28.7 | 107 | 27.2 | 38.2 |
| *MDMA* | 45.4 | 30.9 | 99 | 39.2 | 51.6 |
| *MDMA polyuse* | 40.1 | 30.8 | 108 | 34.2 | 46.0 |
| *Stim* | 32.6 | 30.1 | 58 | 24.6 | 40.6 |
| *Can/Alc* | 29.4 | 25.0 | 68 | 23.3 | 35.5 |
| *No-Use* | 37.5 | 30.0 | 216 | 33.5 | 41.5 |
| *All* | 36.7 | 29.7 | 740 | 34.6 | 38.9 |

*Feeling of Guilt*

| *Substances group* | *Mean* | *SD* | *N* | *Lower CI* | *Upper CI* |
|---|---|---|---|---|---|
| *Halluc* | 42.0 | 31.5 | 84 | 35.1 | 48.8 |
| *Halluc polyuse* | 43.4 | 34.9 | 107 | 36.6 | 50.1 |
| *MDMA* | 40.7 | 34.1 | 99 | 33.9 | 47.6 |
| *MDMA polyuse* | 39.8 | 34.2 | 108 | 33.2 | 46.4 |
| *Stim* | 47.2 | 31.6 | 58 | 38.8 | 55.6 |
| *Can/Alc* | 47.1 | 32.9 | 68 | 39.1 | 55.1 |
| *No-Use* | 39.4 | 32.7 | 216 | 35.0 | 43.8 |
| *All* | 41.8 | 33.4 | 740 | 39.4 | 44.2 |

*Feeling Overwhelmed*

| *Substances group* | *Mean* | *SD* | *N* | *Lower CI* | *Upper CI* |
|---|---|---|---|---|---|
| *Halluc* | 74.0 | 24.6 | 84 | 68.7 | 79.4 |
| *Halluc polyuse* | 77.2 | 21.2 | 107 | 73.1 | 81.3 |
| *MDMA* | 66.6 | 25.2 | 99 | 61.5 | 71.6 |
| *MDMA polyuse* | 74.7 | 20.7 | 108 | 70.7 | 78.6 |
| *Stim* | 79.0 | 18.9 | 58 | 74.0 | 84.0 |
| *Can/Alc* | 79.7 | 21.3 | 68 | 74.6 | 84.9 |
| *No-Use* | 73.5 | 23.8 | 216 | 70.3 | 76.7 |
| *All* | 74.4 | 23.0 | 740 | 72.7 | 76.0 |

*Mental Distress (K6 score)*

| *Substances group* | *Mean* | *SD* | *N* | *Lower CI* | *Upper CI* |
|---|---|---|---|---|---|
| *Halluc* | 12.5 | 5.9 | 54 | 10.9 | 14.2 |
| *Halluc polyuse* | 12.5 | 5.2 | 75 | 11.3 | 13.7 |
| *MDMA* | 10.5 | 5.1 | 63 | 9.2 | 11.8 |
| *MDMA polyuse* | 11.4 | 5.2 | 71 | 10.2 | 12.7 |
| *Stim* | 13.3 | 5.2 | 34 | 11.4 | 15.1 |
| *Can/Alc* | 14.4 | 4.1 | 33 | 12.9 | 15.9 |
| *No-Use* | 12.2 | 5.1 | 104 | 11.2 | 13.2 |
| *All* | 12.2 | 5.3 | 434 | 11.7 | 12.7 |

*PTSD symptoms severity (PCL-5 scores)*

| *Substances group* | *Mean* | *SD* | *N* | *Lower CI* | *Upper CI* |
|---|---|---|---|---|---|

| | | | | | |
|---|---|---|---|---|---|
| *Halluc* | 44.5 | 15.5 | 48 | 39.9 | 49.1 |
| *Halluc polyuse* | 41.7 | 15.2 | 74 | 38.2 | 45.3 |
| *MDMA* | 37.3 | 15.2 | 59 | 33.3 | 41.2 |
| *MDMA polyuse* | 38.9 | 16.5 | 68 | 34.8 | 42.9 |
| *Stim* | 43.3 | 11.7 | 33 | 39.1 | 47.5 |
| *Can/Alc* | 48.3 | 12.8 | 32 | 43.6 | 53.0 |
| *No-Use* | 39.8 | 14.9 | 94 | 36.7 | 42.9 |
| *All* | 41.1 | 15.3 | 408 | 39.6 | 42.6 |

*Main Analysis*

For each measure evaluated in the main analysis (including the groups: *No-Use*, *Can/Alc*, *MDMA*, *Halluc*), we present the results for two linear regression models:

- Model 1: The first model comparing individuals in the *No-Use* group to each of the other groups.
- Model 2: The second model, adjusted for age, sex, and days since the TE, comparing the *No-Use* group to the other groups.

Note: One exception is made for the "Helpfulness" measure. Since no a priori hypothesis was established, we compared each group against all others. The *Can/Alc* group is used as the reference for the following reasons: (1) the *No-Use* group did not answer this question, as it was not relevant to them; (2) the *Can/Alc* group had significantly lower helpfulness ratings than all other groups; and (3) no significant differences were found among the remaining groups. For brevity, we present two models using the *Can/Alc* group as the reference.

**Model 1:**

$$\boldsymbol{Control = X_0 + Substances}$$

Reference group: *No-Use*

| | |
|---|---|
| ***No. Observations*** | 467 |
| ***DoF Residuals*** | 463 |
| ***DoF Model*** | 3 |
| ***R-squared*** | 0.004 |
| ***Adj. R-squared*** | -0.003 |
| ***F-statistic*** | 0.5874 |
| ***Prob (F-statistic)*** | 0.624 |
| ***BIC*** | 4674 |

| | *Coefficient* | *SE* | *t* | *p* | *Lower CI* | *Upper CI* |
|---|---|---|---|---|---|---|
| *Intercept* | 35.5 | 2.4 | 14.8 | 0.0 | 30.8 | 40.3 |
| *Halluc* | 0.1 | 4.5 | 0.0 | 0.981 | -8.8 | 9.0 |
| *MDMA* | 1.3 | 4.3 | 0.3 | 0.765 | -7.1 | 9.7 |
| *Can/Alc* | -5.6 | 4.9 | -1.1 | 0.251 | -15.3 | 4.0 |

**Model 2:**

$$\boldsymbol{Control = X_0 + Substances + Age + Sex + Days\ from\ event}$$

Reference group: *No-Use*

| | |
|---|---|
| ***No. Observations*** | 467 |
| ***DoF Residuals*** | 460 |
| ***DoF Model*** | 6 |
| ***R-squared*** | 0.024 |
| ***Adj. R-squared*** | 0.011 |
| ***F-statistic*** | 1.866 |
| ***Prob (F-statistic)*** | 0.085 |
| ***BIC*** | 4683 |

| | *Coefficient* | *SE* | *t* | *p* | *Lower CI* | *Upper CI* |
|---|---|---|---|---|---|---|
| *Intercept* | 40.7 | 7.8 | 5.2 | 0.0 | 25.3 | 56.1 |
| *Halluc* | -1.5 | 4.6 | -0.3 | 0.735 | -10.5 | 7.4 |
| *MDMA* | -0.5 | 4.3 | -0.1 | 0.911 | -9.0 | 8.0 |
| *Can/Alc* | -6.6 | 4.9 | -1.3 | 0.18 | -16.2 | 3.0 |
| *Age* | 0.0 | 0.2 | 0.0 | 0.979 | -0.5 | 0.4 |
| *Sex* | 7.0 | 3.4 | 2.1 | 0.039 | 0.4 | 13.7 |
| *from TE Days* | -0.1 | 0.1 | -2.4 | 0.018 | -0.2 | 0.0 |

**Model 1:**

$$\boldsymbol{Isolation = X_0 + Substances}$$

Reference group: *No-Use*

| | |
|---|---|
| ***No. Observations*** | 467 |
| ***DoF Residuals*** | 463 |
| ***DoF Model*** | 3 |
| ***R-squared*** | 0.010 |
| ***Adj. R-squared*** | 0.003 |
| ***F-statistic*** | 1.526 |
| ***Prob (F-statistic)*** | 0.207 |
| ***BIC*** | 4702 |

| | *Coefficient* | *SE* | *t* | *p* | *Lower CI* | *Upper CI* |
|---|---|---|---|---|---|---|
| *Intercept* | 47.3 | 2.5 | 19.1 | 0.0 | 42.4 | 52.1 |

| | | | | | | |
|---|---|---|---|---|---|---|
| *Halluc* | -1.2 | 4.7 | -0.3 | 0.795 | -10.4 | 8.0 |
| *MDMA* | -8.8 | 4.4 | -2.0 | 0.046 | -17.5 | -0.2 |
| *Can/Alc* | 0.8 | 5.1 | 0.2 | 0.881 | -9.2 | 10.7 |

**Model 2:**

$$Isolation = X_0 + Substances + Age + Sex + Days\ from\ event$$

Reference group: *No-Use*

| | |
|---|---|
| ***No. Observations*** | 467 |
| ***DoF Residuals*** | 460 |
| ***DoF Model*** | 6 |
| ***R-squared*** | 0.012 |
| ***Adj. R-squared*** | -0.001 |
| ***F-statistic*** | 0.8971 |
| ***Prob (F-statistic)*** | 0.497 |
| ***BIC*** | 4720 |

| | *Coefficient* | *SE* | *t* | *p* | *Lower CI* | *Upper CI* |
|---|---|---|---|---|---|---|
| *Intercept* | 50.6 | 8.2 | 6.2 | 0.0 | 34.5 | 66.6 |
| *Halluc* | -1.8 | 4.8 | -0.4 | 0.712 | -11.1 | 7.6 |
| *MDMA* | -9.6 | 4.5 | -2.1 | 0.033 | -18.5 | -0.8 |
| *Can/Alc* | 0.5 | 5.1 | 0.1 | 0.916 | -9.4 | 10.5 |
| *Age* | -0.1 | 0.2 | -0.5 | 0.593 | -0.6 | 0.3 |
| *Sex* | 2.3 | 3.5 | 0.7 | 0.506 | -4.6 | 9.3 |
| *Days from TE* | 0.0 | 0.1 | -0.2 | 0.812 | -0.1 | 0.1 |

**Model 1:** $Helpfulness = X_0 + Substances$

Reference group: *Can/Alc*

| | |
|---|---|
| ***No. Observations*** | 250 |
| ***DoF Residuals*** | 247 |
| ***DoF Model*** | 2 |
| ***R-squared*** | 0.047 |
| ***Adj. R-squared*** | 0.040 |
| ***F-statistic*** | 6.144 |
| ***Prob (F-statistic)*** | 0.00249 |
| ***BIC*** | 2312 |

| | *Coefficient* | *SE* | *t* | *p* | *Lower CI* | *Upper CI* |
|---|---|---|---|---|---|---|
| *Intercept* | 50.2 | 2.9 | 17.3 | 0.0 | 44.5 | 55.9 |
| *Halluc* | 11.3 | 3.9 | 2.9 | 0.001 | 3.6 | 19.0 |
| *MDMA* | 12.4 | 3.8 | 3.3 | 0.004 | 4.9 | 19.9 |

**Model 2:**

$$\boldsymbol{Helpfulness = X_0 + Substances + Age + Sex + Days\ from\ event}$$

Reference group: *Can/Alc*

| | |
|---|---|
| ***No. Observations*** | 250 |
| ***DoF Residuals*** | 244 |
| ***DoF Model*** | 5 |
| ***R-squared*** | 0.054 |
| ***Adj. R-squared*** | 0.034 |
| ***F-statistic*** | 2.779 |
| ***Prob (F-statistic)*** | 0.0184 |
| ***BIC*** | 2327 |

| | *Coefficient* | *SE* | *t* | *p* | *Lower CI* | *Upper CI* |
|---|---|---|---|---|---|---|
| *Intercept* | 50.3 | 8.5 | 5.9 | 0.0 | 33.6 | 67.0 |
| *Halluc* | 11.5 | 4.0 | 2.9 | 0.004 | 3.6 | 19.3 |
| *MDMA* | 12.4 | 3.9 | 3.2 | 0.001 | 4.8 | 20.0 |
| *Age* | -0.1 | 0.2 | -0.4 | 0.697 | -0.6 | 0.4 |
| *Sex* | -1.8 | 3.3 | -0.5 | 0.586 | -8.3 | 4.7 |
| *Days from TE* | 0.1 | 0.1 | 1.1 | 0.265 | 0.0 | 0.2 |

**Model 1:**

$$\boldsymbol{Interactions = X_0 + Substances}$$

Reference group: *No-Use*

| | |
|---|---|
| ***No. Observations*** | 467 |
| ***DoF Residuals*** | 463 |
| ***DoF Model*** | 3 |
| ***R-squared*** | 0.028 |
| ***Adj. R-squared*** | 0.022 |
| ***F-statistic*** | 4.493 |
| ***Prob (F-statistic)*** | 0.00403 |

| | |
|---|---|
| ***BIC*** | 4449 |

| | *Coefficient* | *SE* | *t* | *p* | *Lower CI* | *Upper CI* |
|---|---|---|---|---|---|---|
| *Intercept* | 66.5 | 1.9 | 35.2 | 0.0 | 62.8 | 70.2 |
| *Halluc* | -3.0 | 3.6 | -0.8 | 0.405 | -10.0 | 4.0 |
| *MDMA* | 10.0 | 3.4 | 3.0 | 0.003 | 3.4 | 16.7 |
| *Can/Alc* | -2.3 | 3.9 | -0.6 | 0.543 | -9.9 | 5.2 |

**Model 2:**

$$\boldsymbol{Interactions = X_0 + Substances + Age + Sex + Days\ from\ event}$$

Reference group: *No-Use*

| | |
|---|---|
| ***No. Observations*** | 467 |
| ***DoF Residuals*** | 460 |
| ***DoF Model*** | 6 |
| ***R-squared*** | 0.054 |
| ***Adj. R-squared*** | 0.042 |
| ***F-statistic*** | 4.381 |
| ***Prob (F-statistic)*** | 0.000256 |
| ***BIC*** | 4455 |

| | *Coefficient* | *SE* | *t* | *p* | *Lower CI* | *Upper CI* |
|---|---|---|---|---|---|---|
| *Intercept* | 84.4 | 6.1 | 13.8 | 0.0 | 72.4 | 96.5 |
| *Halluc* | -3.5 | 3.6 | -1.0 | 0.331 | -10.5 | 3.6 |
| *MDMA* | 8.2 | 3.4 | 2.4 | 0.016 | 1.5 | 14.9 |
| *Can/Alc* | -2.8 | 3.8 | -0.7 | 0.458 | -10.4 | 4.7 |
| *Age* | -0.5 | 0.2 | -2.7 | 0.008 | -0.8 | -0.1 |
| *Sex* | 2.3 | 2.7 | 0.9 | 0.385 | -2.9 | 7.5 |
| *Days from TE* | -0.1 | 0.0 | -2.0 | 0.044 | -0.2 | 0.0 |

**Model 1:**

$$\boldsymbol{Support = X_0 + Substances}$$

Reference group: *No-Use*

| | |
|---|---|
| ***No. Observations*** | 467 |
| ***DoF Residuals*** | 463 |

| | |
|---|---|
| ***DoF Model*** | 3 |
| ***R-squared*** | 0.016 |
| ***Adj. R-squared*** | 0.010 |
| ***F-statistic*** | 2.527 |
| ***Prob (F-statistic)*** | 0.0568 |
| ***BIC*** | 4346 |

| | *Coefficient* | *SE* | *t* | *p* | *Lower CI* | *Upper CI* |
|---|---|---|---|---|---|---|
| *Intercept* | 76.3 | 1.7 | 45.2 | 0.0 | 73.0 | 79.6 |
| *Halluc* | -3.7 | 3.2 | -1.2 | 0.245 | -10.0 | 2.6 |
| *MDMA* | 3.7 | 3.0 | 1.2 | 0.221 | -2.2 | 9.6 |
| *Can/Alc* | -6.0 | 3.5 | -1.7 | 0.084 | -12.8 | 0.8 |

**Model 2:**

$$\boldsymbol{Support = X_0 + Substances + Age + Sex + Days\ from\ event}$$

Reference group: *No-Use*

| | |
|---|---|
| ***No. Observations*** | 467 |
| ***DoF Residuals*** | 460 |
| ***DoF Model*** | 6 |
| ***R-squared*** | 0.034 |
| ***Adj. R-squared*** | 0.021 |
| ***F-statistic*** | 2.667 |
| ***Prob (F-statistic)*** | 0.0149 |
| ***BIC*** | 4356 |

| | *Coefficient* | *SE* | *t* | *p* | *Lower CI* | *Upper CI* |
|---|---|---|---|---|---|---|
| *Intercept* | 86.2 | 5.5 | 15.6 | 0.0 | 75.3 | 97.0 |
| *Halluc* | -4.2 | 3.2 | -1.3 | 0.192 | -10.5 | 2.1 |
| *MDMA* | 2.6 | 3.1 | 0.9 | 0.389 | -3.4 | 8.6 |
| *Can/Alc* | -6.5 | 3.4 | -1.9 | 0.061 | -13.2 | 0.3 |
| *Age* | -0.2 | 0.2 | -1.0 | 0.32 | -0.5 | 0.2 |
| *Sex* | 2.1 | 2.4 | 0.9 | 0.37 | -2.6 | 6.8 |
| *Days from TE* | -0.1 | 0.0 | -2.5 | 0.012 | -0.2 | 0.0 |

**Model 1:**

$$\boldsymbol{Sleep = X_0 + Substances}$$

Reference group: *No-Use*

| | |
|---|---|
| ***No. Observations*** | 467 |
| ***DoF Residuals*** | 463 |
| ***DoF Model*** | 3 |
| ***R-squared*** | 0.029 |
| ***Adj. R-squared*** | 0.023 |
| ***F-statistic*** | 4.595 |
| ***Prob (F-statistic)*** | 0.0035 |
| ***BIC*** | 4497 |

| | *Coefficient* | *SE* | *t* | *p* | *Lower CI* | *Upper CI* |
|---|---|---|---|---|---|---|
| *Intercept* | 37.5 | 2.0 | 18.9 | 0.0 | 33.6 | 41.4 |
| *Halluc* | -3.5 | 3.8 | -0.9 | 0.351 | -10.9 | 3.9 |
| *MDMA* | 7.9 | 3.5 | 2.2 | 0.025 | 1.0 | 14.9 |
| *Can/Alc* | -8.1 | 4.1 | -2.0 | 0.046 | -16.1 | -0.1 |

**Model 2:**

$$\boldsymbol{Sleep = X_0 + Substances + Age + Sex + Days\ from\ event}$$

Reference group: *No-Use*

| | |
|---|---|
| ***No. Observations*** | 467 |
| ***DoF Residuals*** | 460 |
| ***DoF Model*** | 6 |
| ***R-squared*** | 0.034 |
| ***Adj. R-squared*** | 0.021 |
| ***F-statistic*** | 2.672 |
| ***Prob (F-statistic)*** | 0.0147 |
| ***BIC*** | 4513 |

| | *Coefficient* | *SE* | *t* | *p* | *Lower CI* | *Upper CI* |
|---|---|---|---|---|---|---|
| *Intercept* | 30.1 | 6.5 | 4.6 | 0.0 | 17.2 | 42.9 |
| *Halluc* | -4.0 | 3.8 | -1.1 | 0.293 | -11.5 | 3.5 |
| *MDMA* | 8.1 | 3.6 | 2.2 | 0.025 | 1.0 | 15.2 |
| *Can/Alc* | -8.3 | 4.1 | -2.0 | 0.043 | -16.3 | -0.3 |
| *Age* | 0.3 | 0.2 | 1.3 | 0.18 | -0.1 | 0.6 |
| *Sex* | 2.1 | 2.8 | 0.7 | 0.457 | -3.5 | 7.7 |

| | | | | | | |
|---|---|---|---|---|---|---|
| *Days from TE* | 0.0 | 0.0 | -0.3 | 0.775 | -0.1 | 0.1 |

**Model 1:**

$$\boldsymbol{Guilt = X_0 + Substances}$$

Reference group: *No-Use*

| | |
|---|---|
| ***No. Observations*** | 467 |
| ***DoF Residuals*** | 463 |
| ***DoF Model*** | 3 |
| ***R-squared*** | 0.006 |
| ***Adj. R-squared*** | -0.000 |
| ***F-statistic*** | 0.9721 |
| ***Prob (F-statistic)*** | 0.406 |
| ***BIC*** | 4610 |

| | *Coefficient* | *SE* | *t* | *p* | *Lower CI* | *Upper CI* |
|---|---|---|---|---|---|---|
| *Intercept* | 39.4 | 2.2 | 17.6 | 0.0 | 35.0 | 43.8 |
| *Halluc* | 2.6 | 4.2 | 0.6 | 0.542 | -5.7 | 10.9 |
| *MDMA* | 1.3 | 4.0 | 0.3 | 0.738 | -6.5 | 9.2 |
| *Can/Alc* | 7.7 | 4.6 | 1.7 | 0.092 | -1.3 | 16.7 |

**Model 2:**

$$\boldsymbol{Guilt = X_0 + Substances + Age + Sex + Days\ from\ event}$$

Reference group: *No-Use*

| | |
|---|---|
| ***No. Observations*** | 467 |
| ***DoF Residuals*** | 460 |
| ***DoF Model*** | 6 |
| ***R-squared*** | 0.034 |
| ***Adj. R-squared*** | 0.021 |
| ***F-statistic*** | 2.660 |
| ***Prob (F-statistic)*** | 0.0151 |
| ***BIC*** | 4616 |

| | *Coefficient* | *SE* | *t* | *p* | *Lower CI* | *Upper CI* |
|---|---|---|---|---|---|---|
| *Intercept* | 43.4 | 7.3 | 6.0 | 0.0 | 29.1 | 57.8 |

| | | | | | | |
|---|---|---|---|---|---|---|
| *Halluc* | 4.9 | 4.3 | 1.2 | 0.251 | -3.5 | 13.3 |
| *MDMA* | 3.5 | 4.0 | 0.9 | 0.387 | -4.4 | 11.4 |
| *Can/Alc* | 8.2 | 4.5 | 1.8 | 0.07 | -0.7 | 17.2 |
| *Age* | 0.2 | 0.2 | 0.8 | 0.4 | -0.2 | 0.6 |
| *Sex* | -10.0 | 3.2 | -3.2 | 0.002 | -16.2 | -3.8 |
| *Days from TE* | -0.1 | 0.1 | -1.2 | 0.224 | -0.2 | 0.0 |

**Model 1:**

$$\boldsymbol{Overwhelmed = X_0 + Substances}$$

Reference group: *No-Use*

| | |
|---|---|
| ***No. Observations*** | 467 |
| ***DoF Residuals*** | 463 |
| ***DoF Model*** | 3 |
| ***R-squared*** | 0.027 |
| ***Adj. R-squared*** | 0.020 |
| ***F-statistic*** | 4.220 |
| ***Prob (F-statistic)*** | 0.00584 |
| ***BIC*** | 4315 |

| | *Coefficient* | *SE* | *t* | *p* | *Lower CI* | *Upper CI* |
|---|---|---|---|---|---|---|
| *Intercept* | 73.5 | 1.6 | 45.0 | 0.0 | 70.3 | 76.8 |
| *Halluc* | 0.5 | 3.1 | 0.2 | 0.873 | -5.6 | 6.6 |
| *MDMA* | -7.0 | 2.9 | -2.4 | 0.017 | -12.7 | -1.2 |
| *Can/Alc* | 6.2 | 3.3 | 1.9 | 0.064 | -0.4 | 12.8 |

**Model 2:**

$$\boldsymbol{Overwhelmed = X_0 + Substances + Age + Sex + Days\ from\ event}$$

Reference group: *No-Use*

| | |
|---|---|
| ***No. Observations*** | 467 |
| ***DoF Residuals*** | 460 |
| ***DoF Model*** | 6 |
| ***R-squared*** | 0.051 |
| ***Adj. R-squared*** | 0.039 |
| ***F-statistic*** | 4.138 |
| ***Prob (F-statistic)*** | 0.000462 |

| | |
|---|---|
| ***BIC*** | 4322 |

| | *Coefficient* | *SE* | *t* | *p* | *Lower CI* | *Upper CI* |
|---|---|---|---|---|---|---|
| *Intercept* | 69.1 | 5.3 | 13.0 | 0.0 | 58.6 | 79.5 |
| *Halluc* | 2.0 | 3.1 | 0.6 | 0.518 | -4.1 | 8.1 |
| *MDMA* | -5.0 | 2.9 | -1.7 | 0.088 | -10.8 | 0.7 |
| *Can/Alc* | 6.7 | 3.3 | 2.0 | 0.045 | 0.2 | 13.2 |
| *Age* | 0.3 | 0.2 | 1.8 | 0.079 | 0.0 | 0.6 |
| *Sex* | -6.6 | 2.3 | -2.8 | 0.005 | -11.1 | -2.0 |
| *Days from TE* | 0.0 | 0.0 | 0.0 | 0.978 | -0.1 | 0.1 |

**Model 1:**

$$K6 = X_0 + Substances$$

Reference group: *No-Use*

| | |
|---|---|
| ***No. Observations*** | 254 |
| ***DoF Residuals*** | 250 |
| ***DoF Model*** | 3 |
| ***R-squared*** | 0.048 |
| ***Adj. R-squared*** | 0.037 |
| ***F-statistic*** | 4.246 |
| ***Prob (F-statistic)*** | 0.00600 |
| ***BIC*** | 1575 |

| | *Coefficient* | *SE* | *t* | *p* | *Lower CI* | *Upper CI* |
|---|---|---|---|---|---|---|
| *Intercept* | 12.2 | 0.5 | 24.0 | 0 | 11.2 | 13.2 |
| *Halluc* | 0.4 | 0. 9 | 0.4 | 0.676 | -1.4 | 2.1 |
| *MDMA* | -1.6 | 0.8 | -2.0 | 0.049 | -3. 3 | -0.0 |
| *Can/Alc* | 2.3 | 1.0 | 2.2 | 0.031 | 0.2 | 4.3 |

**Model 2:**

$$k6 = X_0 + Substances + Age + Sex + Days\ from\ event$$

Reference group: *No-Use*

| | |
|---|---|
| ***No. Observations*** | 254 |
| ***DoF Residuals*** | 247 |
| ***DoF Model*** | 6 |

| | |
|---|---|
| ***R-squared*** | 0.104 |
| ***Adj. R-squared*** | 0.083 |
| ***F-statistic*** | 4.797 |
| ***Prob (F-statistic)*** | 0.00012 |
| ***BIC*** | 1576 |

| | *Coefficient* | *SE* | *t* | *p* | *Lower CI* | *Upper CI* |
|---|---|---|---|---|---|---|
| *Intercept* | 10.7 | 1.5 | 7.0 | 0.0 | 7.7 | 13.7 |
| *Halluc* | 0.8 | 0.9 | 0.9 | 0.372 | -0.9 | 2.5 |
| *MDMA* | -1.2 | 0.8 | -1.5 | 0.147 | -2.8 | 0.4 |
| *Can/Alc* | 2.1 | 1.0 | 2.1 | 0.038 | 0.1 | 4.1 |
| *Age* | 0.1 | 0.0 | 1.2 | 0.233 | 0.0 | 0.1 |
| *Sex* | -2.2 | 0.7 | -3.3 | 0.001 | -3.5 | -0.9 |
| *Days from TE* | 0.0 | 0.0 | 1.5 | 0.134 | 0.0 | 0.0 |

**Model 1:**

$$PCL = X_0 + Substances$$

Reference group: *No-Use*

| | |
|---|---|
| ***No. Observations*** | 233 |
| ***DoF Residuals*** | 229 |
| ***DoF Model*** | 3 |
| ***R-squared*** | 0.059 |
| ***Adj. R-squared*** | 0.047 |
| ***F-statistic*** | 4.791 |
| ***Prob (F-statistic)*** | 0.00295 |
| ***BIC*** | 1940 |

| | *Coefficient* | *SE* | *t* | *p* | *Lower CI* | *Upper CI* |
|---|---|---|---|---|---|---|
| *Intercept* | 39.8 | 1.5 | 25.8 | 0.0 | 36.8 | 42.9 |
| *Halluc* | 4.7 | 2.7 | 1.8 | 0.079 | -0.6 | 9.9 |
| *MDMA* | -2.6 | 2.5 | -1.0 | 0.304 | -7.5 | 2.3 |
| *Can/Alc* | 8.5 | 3.1 | 2.8 | 0.006 | 2.4 | 14.5 |

**Model 2:**

$$PCL = X_0 + Substances + Age + Sex + Days\ from\ event$$

Reference group: *No-Use*

| | |
|---|---|
| ***No. Observations*** | 233 |
| ***DoF Residuals*** | 226 |
| ***DoF Model*** | 6 |
| ***R-squared*** | 0.107 |
| ***Adj. R-squared*** | 0.083 |
| ***F-statistic*** | 4.492 |
| ***Prob (F-statistic)*** | 0.000255 |
| ***BIC*** | 1944 |

| | *Coefficient* | *SE* | *t* | *p* | *Lower CI* | *Upper CI* |
|---|---|---|---|---|---|---|
| *Intercept* | 35.4 | 4.5 | 7.8 | 0.0 | 26.4 | 44.4 |
| *Halluc* | 5.8 | 2.6 | 2.2 | 0.029 | 0.6 | 11.0 |
| *MDMA* | -1.2 | 2.5 | -0.5 | 0.637 | -6.1 | 3.7 |
| *Can/Alc* | 7.9 | 3.0 | 2.6 | 0.009 | 2.0 | 13.9 |
| *Age* | 0.2 | 0.1 | 1.4 | 0.173 | -0.1 | 0.4 |
| *Sex* | -5.8 | 2.0 | -2.9 | 0.004 | -9.7 | -1.9 |
| *Days from TE* | 0.0 | 0.0 | 1.0 | 0.331 | 0.0 | 0.1 |

The analyses for the constructs: "Feeling of Control," "Feeling of Isolation," and "Feeling of Support" did not yield significant results. Corresponding distribution graphs for these variables are provided in Figure S1.

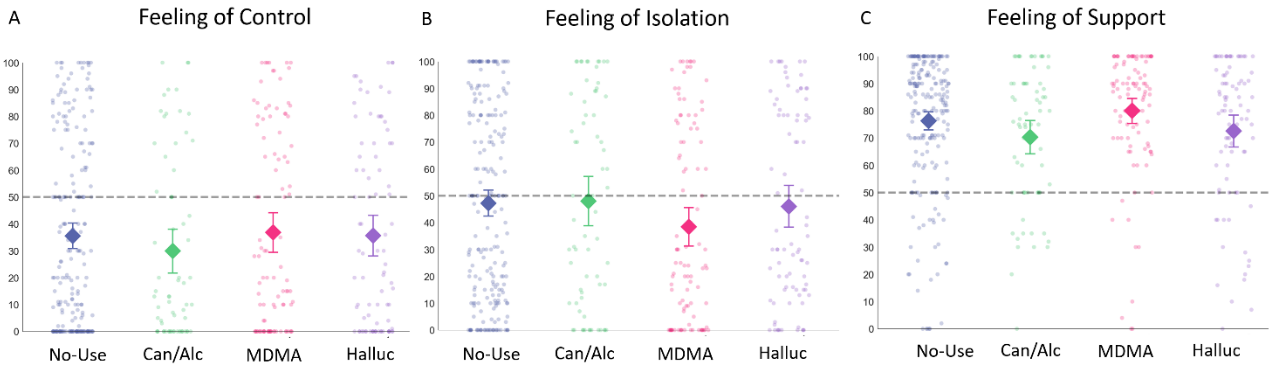


**Figure S1: Outcome Measures Across Groups for the Main Analysis.** Panels A-C represent dot plots with overlaying mean values (diamonds) and 95% confidence intervals (error bars) for the respective outcome measures in relation to substance use categories (*Halluc*, *MDMA*, *Can/Alc*, *No-Use*). **(A) Feelings of Control.** Dots represent participants' self-reported feelings of control during the TE on a scale between 0-100, as higher ratings represent more feelings of control. No significant differences were found between groups in feelings of control during the TE ($F(3,463)=0.59, p=0.62$). **(B) Feelings of Isolation.** Dots represent participants' self-reported feelings of isolation during the TE on a scale between 0-100, as higher ratings represent more feelings of isolation. No significant differences were found between groups in judgments regarding feelings of isolation during the TE ($F(3,643)=1.53, p=0.21$). **(C) Feelings of Support.** Dots represent participants' self-reported feelings of support from friends and family in the peritraumatic period on a scale between 0-100, as higher ratings represent feeling more supported. No significant differences were found between groups in judgments regarding feelings of support during the peritraumatic processing phase ($F(3,463)=2.5, p=0.056$).

*Comprehensive Analysis*

For each measure we also performed a comprehensive analysis including all groups (*No-Use, Can/Alc*, *Stim*, *MDMA*, *MDMA polyuse Halluc*, *Halluc polyuse*), we present the results for two linear regression models:

- Model 1: The first model comparing individuals in the *No-Use* group to each of the other groups.
- Model 2: The second model, adjusted for age, sex, and days since the TE, comparing the *No-Use* group to the other groups.

The findings are depicted in Figure S2.

**Model 1:** $\boldsymbol{Control = X_0 + Substances}$

Reference group: *No-Use*

| | |
|---|---|
| ***No. Observations*** | 740 |
| ***DoF Residuals*** | 733 |
| ***DoF Model*** | 6 |
| ***R-squared*** | 0.005 |
| ***Adj. R-squared*** | -0.003 |
| ***F-statistic*** | 0.5707 |
| ***Prob (F-statistic)*** | 0.754 |
| ***BIC*** | 7407 |

| | *Coefficient* | *SE* | *t* | *p* | *Lower CI* | *Upper CI* |
|---|---|---|---|---|---|---|
| *Intercept* | 35.5 | 2.4 | 14.9 | 0.0 | 30.8 | 40.2 |
| *Halluc* | 0.1 | 4.5 | 0.0 | 1.0 | -8.8 | 9.0 |
| *Halluc polyuse* | 0.7 | 4.2 | 0.2 | 0.9 | -7.4 | 8.9 |
| *MDMA* | 1.3 | 4.3 | 0.3 | 0.8 | -7.1 | 9.7 |
| *MDMA polyuse* | 2.4 | 4.1 | 0.6 | 0.6 | -5.7 | 10.6 |
| *Stim* | -4.6 | 5.2 | -0.9 | 0.4 | -14.8 | 5.6 |
| *Can/Alc* | -5.6 | 4.9 | -1.2 | 0.2 | -15.2 | 4.0 |

**Model 2:** $\boldsymbol{Control = X_0 + Substances + Age + Sex + Days\ from\ event}$

Reference group: *No-Use*

| | |
|---|---|
| ***No. Observations*** | 740 |
| ***DoF Residuals*** | 730 |
| ***DoF Model*** | 9 |
| ***R-squared*** | 0.019 |
| ***Adj. R-squared*** | 0.007 |
| ***F-statistic*** | 1.598 |
| ***Prob (F-statistic)*** | 0.112 |
| ***BIC*** | 7416 |

| | *Coefficient* | *SE* | *t* | *p* | *Lower CI* | *Upper CI* |
|---|---|---|---|---|---|---|
| *Intercept* | 45.0 | 6.8 | 6.6 | 0.0 | 31.6 | 58.4 |
| *Halluc* | -1.1 | 4.5 | -0.2 | 0.8 | -10.0 | 7.8 |
| *Halluc polyuse* | -1.3 | 4.2 | -0.3 | 0.8 | -9.6 | 7.0 |
| *MDMA* | -0.4 | 4.3 | -0.1 | 0.9 | -8.9 | 8.0 |
| *MDMA polyuse* | 0.4 | 4.2 | 0.1 | 0.9 | -7.9 | 8.6 |
| *Stim* | -4.5 | 5.2 | -0.9 | 0.4 | -14.7 | 5.7 |
| *Can/Alc* | -6.4 | 4.9 | -1.3 | 0.2 | -16.0 | 3.2 |
| *Age* | -0.2 | 0.2 | -0.8 | 0.4 | -0.5 | 0.2 |
| *Sex* | 5.1 | 2.7 | 1.9 | 0.1 | -0.3 | 10.5 |
| *Days from TE* | -0.1 | 0.05 | -2.6 | 0.008 | -0.2 | -0.03 |

**Model 1:** $\boldsymbol{Isolation = X_0 + Substances}$

Reference group: *No-Use*

| | |
|---|---|
| ***No. Observations*** | 740 |
| ***DoF Residuals*** | 733 |
| ***DoF Model*** | 6 |
| ***R-squared*** | 0.008 |
| ***Adj. R-squared*** | -0.000 |
| ***F-statistic*** | 0.9535 |
| ***Prob (F-statistic)*** | 0.456 |
| ***BIC*** | 7478 |

| | *Coefficient* | *SE* | *t* | *p* | *Lower CI* | *Upper CI* |
|---|---|---|---|---|---|---|
| *Intercept* | 47.3 | 2.5 | 18.8 | 0.0 | 42.3 | 52.2 |
| *Halluc* | -1.2 | 4.7 | -0.3 | 0.8 | -10.5 | 8.1 |

| | | | | | | |
|---|---|---|---|---|---|---|
| *Halluc polyuse* | 0.7 | 4.4 | 0.2 | 0.9 | -7.8 | 9.3 |
| *MDMA* | -8.8 | 4.5 | -2.0 | 0.049 | -17.6 | -0.04 |
| *MDMA polyuse* | 1.1 | 4.3 | 0.2 | 0.8 | -7.5 | 9.6 |
| *Stim* | 2.0 | 5.5 | 0.4 | 0.7 | -8.7 | 12.7 |
| *Can/Alc* | 0.8 | 5.1 | 0.1 | 0.9 | -9.3 | 10.8 |

**Model 2:** $Isolation = X_0 + Substances + Age + Sex + Days\ from\ event$

Reference group: *No-Use*

| | |
|---|---|
| ***No. Observations*** | 740 |
| ***DoF Residuals*** | 730 |
| ***DoF Model*** | 9 |
| ***R-squared*** | 0.009 |
| ***Adj. R-squared*** | -0.003 |
| ***F-statistic*** | 0.7244 |
| ***Prob (F-statistic)*** | 0.687 |
| ***BIC*** | 7497 |

| | *Coefficient* | *SE* | *t* | *p* | *Lower CI* | *Upper CI* |
|---|---|---|---|---|---|---|
| *Intercept* | 50.8 | 7.2 | 7.1 | 0.0 | 36.6 | 64.9 |
| *Halluc* | -1.6 | 4.8 | -0.3 | 0.7 | -11.0 | 7.8 |
| *Halluc polyuse* | 0.1 | 4.5 | 0.0 | 1.0 | -8.6 | 8.9 |
| *MDMA* | -9.5 | 4.5 | -2.1 | 0.0 | -18.4 | -0.6 |
| *MDMA polyuse* | 0.3 | 4.4 | 0.1 | 0.9 | -8.4 | 9.0 |
| *Stim* | 1.8 | 5.5 | 0.3 | 0.7 | -8.9 | 12.6 |
| *Can/Alc* | 0.6 | 5.1 | 0.1 | 0.9 | -9.5 | 10.7 |
| *Age* | -0.1 | 0.2 | -0.6 | 0.5 | -0.5 | 0.3 |
| *Sex* | 1.6 | 2.9 | 0.6 | 0.6 | -4.0 | 7.3 |
| *Days from TE* | -0.009 | 0.05 | -0.2 | 0.9 | -0.1 | 0.1 |

**Model 1:** $Helpfulness = X_0 + Substances$

Reference group: *Can/Alc*

| | |
|---|---|
| ***No. Observations*** | 522 |
| ***DoF Residuals*** | 516 |
| ***DoF Model*** | 5 |
| ***R-squared*** | 0.026 |
| ***Adj. R-squared*** | 0.017 |

| | |
|---|---|
| ***F-statistic*** | 2.786 |
| ***Prob (F-statistic)*** | 0.017 |
| ***BIC*** | 4839 |

| | *Coefficient* | *SE* | *t* | *p* | *Lower CI* | *Upper CI* |
|---|---|---|---|---|---|---|
| *Intercept* | 50.2 | 2.9 | 17.1 | 0.0 | 44.4 | 56.0 |
| *Halluc* | 11.3 | 3.9 | 2.9 | 0.004 | 3.6 | 19.1 |
| *Halluc polyuse* | 12.3 | 3.8 | 3.3 | 0.001 | 4.9 | 19.7 |
| *MDMA* | 12.4 | 3.8 | 3.2 | 0.001 | 4.9 | 19.9 |
| *MDMA polyuse* | 10.5 | 3.7 | 2.8 | 0.005 | 3.2 | 17.9 |
| *Stim* | 10.8 | 4.3 | 2.5 | 0.013 | 2.3 | 19.3 |

**Model 2:**

$$\boldsymbol{Helpfulness = X_0 + Substances + Age + Sex + Days\ from\ event}$$

Reference group: *Can/Alc*

| | |
|---|---|
| ***No. Observations*** | 522 |
| ***DoF Residuals*** | 513 |
| ***DoF Model*** | 8 |
| ***R-squared*** | 0.033 |
| ***Adj. R-squared*** | 0.018 |
| ***F-statistic*** | 2.172 |
| ***Prob (F-statistic)*** | 0.0282 |
| ***BIC*** | 4854 |

| | *Coefficient* | *SE* | *t* | *p* | *Lower CI* | *Upper CI* |
|---|---|---|---|---|---|---|
| *Intercept* | 49.9 | 6.5 | 7.6 | 0.0 | 37.1 | 62.7 |
| *Halluc* | 11.4 | 4.0 | 2.9 | 0.004 | 3.6 | 19.2 |
| *Halluc polyuse* | 12.6 | 3.8 | 3.3 | 0.001 | 5.2 | 20.1 |
| *MDMA* | 12.4 | 3.8 | 3.2 | 0.001 | 4.8 | 19.9 |
| *MDMA polyuse* | 10.6 | 3.8 | 2.8 | 0.005 | 3.2 | 18.0 |
| *Stim* | 10.2 | 4.3 | 2.3 | 0.02 | 1.6 | 18.7 |
| *Age* | -0.1 | 0.2 | -0.5 | 0.6 | -0.5 | 0.3 |
| *Sex* | -1.6 | 2.3 | -0.7 | 0.5 | -6.1 | 3.0 |
| *Days from TE* | 0.1 | 0.04 | 1.7 | 0.1 | -0.01 | 0.1 |

**Model 1:** $\boldsymbol{Interactions = X_0 + Substances}$

Reference group: *No-Use*

| | |
|---|---|
| ***No. Observations*** | 740 |
| ***DoF Residuals*** | 733 |
| ***DoF Model*** | 6 |
| ***R-squared*** | 0.027 |
| ***Adj. R-squared*** | 0.019 |
| ***F-statistic*** | 3.33 |
| ***Prob (F-statistic)*** | 0.00305 |
| ***BIC*** | 7040 |

| | *Coefficient* | *SE* | *t* | *p* | *Lower CI* | *Upper CI* |
|---|---|---|---|---|---|---|
| *Intercept* | 66.5 | 1.9 | 35.6 | 0.0 | 62.8 | 70.2 |
| *Halluc* | -3.0 | 3.5 | -0.8 | 0.4 | -9.9 | 4.0 |
| *Halluc polyuse* | -1.0 | 3.2 | -0.3 | 0.8 | -7.3 | 5.4 |
| *MDMA* | 10.0 | 3.3 | 3.0 | 0.003 | 3.5 | 16.6 |
| *MDMA polyuse* | 7.7 | 3.2 | 2.4 | 0.0 | 1.3 | 14.0 |
| *Stim* | 2.0 | 4.1 | 0.5 | 0.6 | -6.0 | 10.0 |
| *Can/Alc* | -2.3 | 3.8 | -0.6 | 0.5 | -9.8 | 5.1 |

**Model 2:**

$$\boldsymbol{Interactions = X_0 + Substances + Age + Sex + Days\ from\ event}$$

Reference group: *No-Use*

| | |
|---|---|
| ***No. Observations*** | 740 |
| ***DoF Residuals*** | 730 |
| ***DoF Model*** | 9 |
| ***R-squared*** | 0.044 |
| ***Adj. R-squared*** | 0.032 |
| ***F-statistic*** | 3.738 |
| ***Prob (F-statistic)*** | 0.000131 |
| ***BIC*** | 7047 |

| | *Coefficient* | *SE* | *t* | *p* | *Lower CI* | *Upper CI* |
|---|---|---|---|---|---|---|
| *Intercept* | 82.0 | 5.3 | 15.5 | 0.0 | 71.6 | 92.4 |

| | | | | | | |
|---|---|---|---|---|---|---|
| *Halluc* | -3.4 | 3.5 | -1.0 | 0.3 | -10.4 | 3.5 |
| *Halluc polyuse* | -2.1 | 3.3 | -0.7 | 0.5 | -8.6 | 4.3 |
| *MDMA* | 8.4 | 3.4 | 2.5 | 0.01 | 1.8 | 15.0 |
| *MDMA polyuse* | 5.5 | 3.3 | 1.7 | 0.1 | -0.9 | 12.0 |
| *Stim* | 1.5 | 4.0 | 0.4 | 0.7 | -6.5 | 9.4 |
| *Can/Alc* | -2.8 | 3.8 | -0.7 | 0.5 | -10.2 | 4.7 |
| *Age* | -0.4 | 0.2 | -2.8 | 0.006 | -0.7 | -0.1 |
| *Sex* | 2.1 | 2.1 | 1.0 | 0.3 | -2.1 | 6.3 |
| *Days from TE* | -0.1 | 0.0 | -2.1 | 0.04 | -0.1 | 0.0 |

**Model 1:** $Support = X_0 + Substances$

Reference group: *No-Use*

| | |
|---|---|
| ***No. Observations*** | 740 |
| ***DoF Residuals*** | 733 |
| ***DoF Model*** | 6 |
| ***R-squared*** | 0.032 |
| ***Adj. R-squared*** | 0.024 |
| ***F-statistic*** | 4.036 |
| ***Prob (F-statistic)*** | 0.000547 |
| ***BIC*** | 6870 |

| | *Coefficient* | *SE* | *t* | *p* | *Lower CI* | *Upper CI* |
|---|---|---|---|---|---|---|
| *Intercept* | 76.3 | 1.7 | 45.9 | 0.0 | 73.0 | 79.6 |
| *Halluc* | -3.7 | 3.1 | -1.2 | 0.2 | -9.9 | 2.5 |
| *Halluc polyuse* | -3.3 | 2.9 | -1.2 | 0.3 | -9.0 | 2.4 |
| *MDMA* | 3.7 | 3.0 | 1.2 | 0.2 | -2.1 | 9.5 |
| *MDMA polyuse* | 8.8 | 2.9 | 3.1 | 0.002 | 3.1 | 14.5 |
| *Stim* | 1.5 | 3.6 | 0.4 | 0.7 | -5.6 | 8.6 |
| *Can/Alc* | -6.0 | 3.4 | -1.8 | 0.1 | -12.7 | 0.7 |

**Model 2:** $Support = X_0 + Substances + Age + Sex + Days\ from\ event$

Reference group: *No-Use*

| | |
|---|---|
| ***No. Observations*** | 740 |
| ***DoF Residuals*** | 730 |
| ***DoF Model*** | 9 |
| ***R-squared*** | 0.054 |

| | |
|---|---|
| ***Adj. R-squared*** | 0.042 |
| ***F-statistic*** | 4.622 |
| ***Prob (F-statistic)*** | 5.79e-06 |
| ***BIC*** | 6873 |

| | *Coefficient* | *SE* | *t* | *p* | *Lower CI* | *Upper CI* |
|---|---|---|---|---|---|---|
| *Intercept* | 89.5 | 4.7 | 19.0 | 0.0 | 80.3 | 98.8 |
| *Halluc* | -4.1 | 3.1 | -1.3 | 0.2 | -10.3 | 2.0 |
| *Halluc polyuse* | -4.4 | 2.9 | -1.5 | 0.1 | -10.1 | 1.3 |
| *MDMA* | 2.4 | 3.0 | 0.8 | 0.4 | -3.4 | 8.3 |
| *MDMA polyuse* | 7.0 | 2.9 | 2.4 | 0.02 | 1.3 | 12.7 |
| *Stim* | 1.3 | 3.6 | 0.4 | 0.7 | -5.7 | 8.4 |
| *Can/Alc* | -6.5 | 3.4 | -1.9 | 0.1 | -13.1 | 0.1 |
| *Age* | -0.2 | 0.1 | -1.8 | 0.1 | -0.5 | 0.02 |
| *Sex* | 1.8 | 1.9 | 0.9 | 0.3 | -1.9 | 5.5 |
| *Days from TE* | -0.1 | 0.03 | -3.5 | 0.0 | -0.2 | -0.05 |
| *Days from TE* | *0.01* | *0.04* | *0.3* | *0.8* | *-0.1* | *0.1* |

**Model 1:** $\boldsymbol{Sleep = X_0 + Substances}$

Reference group: *No-Use*

| | |
|---|---|
| ***No. Observations*** | 740 |
| ***DoF Residuals*** | 733 |
| ***DoF Model*** | 6 |
| ***R-squared*** | 0.024 |
| ***Adj. R-squared*** | 0.016 |
| ***F-statistic*** | 3.043 |
| ***Prob (F-statistic)*** | 0.00603 |
| ***BIC*** | 7148 |

| | *Coefficient* | *SE* | *t* | *p* | *Lower CI* | *Upper CI* |
|---|---|---|---|---|---|---|
| *Intercept* | 37.5 | 2.0 | 18.7 | 0.0 | 33.5 | 41.4 |
| *Halluc* | -3.5 | 3.8 | -0.9 | 0.4 | -11.0 | 3.9 |
| *Halluc polyuse* | -4.8 | 3.5 | -1.4 | 0.2 | -11.6 | 2.1 |
| *MDMA* | 7.9 | 3.6 | 2.2 | 0.027 | 0.9 | 15.0 |

| | | | | | | |
|---|---|---|---|---|---|---|
| *MDMA polyuse* | 2.6 | 3.5 | 0.8 | 0.4 | -4.2 | 9.5 |
| *Stim* | -4.9 | 4.4 | -1.1 | 0.3 | -13.4 | 3.7 |
| *Can/Alc* | -8.1 | 4.1 | -2.0 | 0.048 | -16.2 | -0.1 |

**Model 2:** $\boldsymbol{Sleep = X_0 + Substances + Age + Sex + Days\ from\ event}$

Reference group: *No-Use*

| | |
|---|---|
| ***No. Observations*** | 740 |
| ***DoF Residuals*** | 730 |
| ***DoF Model*** | 9 |
| ***R-squared*** | 0.025 |
| ***Adj. R-squared*** | 0.013 |
| ***F-statistic*** | 2.096 |
| ***Prob (F-statistic)*** | 0.0278 |
| ***BIC*** | 7167 |

| | *Coefficient* | *SE* | *t* | *p* | *Lower CI* | *Upper CI* |
|---|---|---|---|---|---|---|
| *Intercept* | 34.2 | 5.8 | 6.0 | 0.0 | 23.0 | 45.5 |
| *Halluc* | -3.9 | 3.8 | -1.0 | 0.3 | -11.4 | 3.7 |
| *Halluc polyuse* | -5.2 | 3.6 | -1.4 | 0.1 | -12.2 | 1.8 |
| *MDMA* | 7.8 | 3.6 | 2.2 | 0.032 | 0.7 | 15.0 |
| *MDMA polyuse* | 2.7 | 3.5 | 0.8 | 0.5 | -4.3 | 9.6 |
| *Stim* | -4.7 | 4.4 | -1.1 | 0.3 | -13.3 | 3.9 |
| *Can/Alc* | -8.2 | 4.1 | -2.0 | 0.047 | -16.3 | -0.1 |
| *Age* | 0.1 | 0.2 | 0.4 | 0.7 | -0.3 | 0.4 |
| *Sex* | 1.5 | 2.3 | 0.7 | 0.5 | -3.0 | 6.1 |
| *Days from TE* | 0.01 | 0.04 | 0.3 | 0.8 | -0.1 | 0.1 |

**Model 1:** $\boldsymbol{Guilt = X_0 + Substances}$

Reference group: *No-Use*

| | |
|---|---|
| ***No. Observations*** | 740 |
| ***DoF Residuals*** | 733 |
| ***DoF Model*** | 6 |
| ***R-squared*** | 0.007 |
| ***Adj. R-squared*** | -0.001 |
| ***F-statistic*** | 0.8549 |
| ***Prob (F-statistic)*** | 0.528 |

| | |
|---|---|
| ***BIC*** | 7332 |

| | *Coefficient* | *SE* | *t* | *p* | *Lower CI* | *Upper CI* |
|---|---|---|---|---|---|---|
| *Intercept* | 39.4 | 2.3 | 17.3 | 0.0 | 34.9 | 43.8 |
| *Halluc* | 2.6 | 4.3 | 0.6 | 0.5 | -5.8 | 11.0 |
| *Halluc polyuse* | 4.0 | 3.9 | 1.0 | 0.3 | -3.8 | 11.7 |
| *MDMA* | 1.3 | 4.1 | 0.3 | 0.7 | -6.6 | 9.3 |
| *MDMA polyuse* | 0.4 | 3.9 | 0.1 | 0.9 | -7.3 | 8.2 |
| *Stim* | 7.9 | 4.9 | 1.6 | 0.1 | -1.8 | 17.6 |
| *Can/Alc* | 7.7 | 4.6 | 1.7 | 0.1 | -1.4 | 16.9 |

**Model 2:** $\boldsymbol{Guilt = X_0 + Substances + Age + Sex + Days\ from\ event}$

Reference group: *No-Use*

| | |
|---|---|
| ***No. Observations*** | 740 |
| ***DoF Residuals*** | 730 |
| ***DoF Model*** | 9 |
| ***R-squared*** | 0.038 |
| ***Adj. R-squared*** | 0.026 |
| ***F-statistic*** | 3.226 |
| ***Prob (F-statistic)*** | 0.000755 |
| ***BIC*** | 7328 |

| | *Coefficient* | *SE* | *t* | *p* | *Lower CI* | *Upper CI* |
|---|---|---|---|---|---|---|
| *Intercept* | 48.3 | 6.4 | 7.5 | 0.0 | 35.7 | 60.9 |
| *Halluc* | 5.3 | 4.3 | 1.2 | 0.2 | -3.1 | 13.7 |
| *Halluc polyuse* | 7.3 | 4.0 | 1.8 | 0.1 | -0.5 | 15.1 |
| *MDMA* | 3.5 | 4.1 | 0.9 | 0.4 | -4.5 | 11.4 |
| *MDMA polyuse* | 1.9 | 4.0 | 0.5 | 0.6 | -5.9 | 9.6 |
| *Stim* | 7.9 | 4.9 | 1.6 | 0.1 | -1.7 | 17.5 |
| *Can/Alc* | 8.3 | 4.6 | 1.8 | 0.1 | -0.7 | 17.3 |
| *Age* | 0.05 | 0.2 | 0.3 | 0.8 | -0.3 | 0.4 |
| *Sex* | -11.6 | 2.6 | -4.5 | 0.0 | -16.7 | -6.5 |
| *Days from TE* | -0.1 | 0.04 | -1.6 | 0.1 | -0.2 | 0.02 |

**Model 1:** $\boldsymbol{Overwhelmed = X_0 + Substances}$

Reference group: *No-Use*

| | |
|---|---|
| ***No. Observations*** | 740 |
| ***DoF Residuals*** | 733 |
| ***DoF Model*** | 6 |
| ***R-squared*** | 0.026 |
| ***Adj. R-squared*** | 0.018 |
| ***F-statistic*** | 3.266 |
| ***Prob (F-statistic)*** | 0.004 |
| ***BIC*** | 6769 |

| | *Coefficient* | *SE* | *t* | *p* | *Lower CI* | *Upper CI* |
|---|---|---|---|---|---|---|
| *Intercept* | 73.5 | 1.6 | 47.3 | 0.0 | 70.5 | 76.6 |
| *Halluc* | 0.5 | 2.9 | 0.2 | 0.9 | -5.3 | 6.3 |
| *Halluc polyuse* | 3.7 | 2.7 | 1.4 | 0.2 | -1.6 | 9.0 |
| *MDMA* | -7.0 | 2.8 | -2.5 | 0.012 | -12.4 | -1.5 |
| *MDMA polyuse* | 1.1 | 2.7 | 0.4 | 0.7 | -4.2 | 6.4 |
| *Stim* | 5.5 | 3.4 | 1.6 | 0.1 | -1.2 | 12.1 |
| *Can/Alc* | 6.2 | 3.2 | 2.0 | 0.052 | -0.04 | 12.4 |

**Model 2:**

$$\boldsymbol{Overwhelmed = X_0 + Substances + Age + Sex + Days\ from\ event}$$

Reference group: *No-Use*

| | |
|---|---|
| ***No. Observations*** | 740 |
| ***DoF Residuals*** | 730 |
| ***DoF Model*** | 9 |
| ***R-squared*** | 0.051 |
| ***Adj. R-squared*** | 0.04 |
| ***F-statistic*** | 4.386 |
| ***Prob (F-statistic)*** | 1.34e-05 |
| ***BIC*** | 6770 |

| | *Coefficient* | *SE* | *t* | *p* | *Lower CI* | *Upper CI* |
|---|---|---|---|---|---|---|

| | | | | | | |
|---|---|---|---|---|---|---|
| *Intercept* | 73.0 | 4.4 | 16.6 | 0.0 | 64.4 | 81.7 |
| *Halluc* | 2.0 | 2.9 | 0.7 | 0.5 | -3.8 | 7.7 |
| *Halluc polyuse* | 5.6 | 2.7 | 2.1 | 0.04 | 0.3 | 11.0 |
| *MDMA* | -5.3 | 2.8 | -1.9 | 0.1 | -10.8 | 0.2 |
| *MDMA polyuse* | 2.5 | 2.7 | 0.9 | 0.3 | -2.8 | 7.9 |
| *Stim* | 5.8 | 3.3 | 1.7 | 0.1 | -0.8 | 12.4 |
| *Can/Alc* | 6.5 | 3.1 | 2.1 | 0.038 | 0.4 | 12.7 |
| *Age* | 0.2 | 0.1 | 1.8 | 0.1 | -0.03 | 0.5 |
| *Sex* | -6.5 | 1.8 | -3.7 | 0.0 | -10.0 | -3.0 |
| *Days from TE* | -0.038 | 0.03 | -1.3 | 0.2 | -0.1 | 0.02 |

**Model 1:** $K6 = X_0 + Substances$

Reference group: *No-Use*

| | |
|---|---|
| ***No. Observations*** | 434 |
| ***DoF Residuals*** | 427 |
| ***DoF Model*** | 6 |
| ***R-squared*** | 0.036 |
| ***Adj. R-squared*** | 0.022 |
| ***F-statistic*** | 2.641 |
| ***Prob (F-statistic)*** | 0.0159 |
| ***BIC*** | 2700 |

| | *Coefficient* | *SE* | *t* | *p* | *Lower CI* | *Upper CI* |
|---|---|---|---|---|---|---|
| *Intercept* | 12.2 | 0.5 | 23.8 | 0.0 | 11.2 | 13.2 |
| *Halluc* | 0.4 | 0.9 | 0.4 | 0.7 | -1.4 | 2.1 |
| *Halluc polyuse* | 0.3 | 0.8 | 0.4 | 0.7 | -1.3 | 1.8 |
| *MDMA* | -1.6 | 0.8 | -2.0 | 0.050 | -3.3 | 0.002 |
| *MDMA polyuse* | -0.7 | 0.8 | -0.9 | 0.4 | -2.3 | 0.8 |
| *Stim* | 1.1 | 1.0 | 1.1 | 0.3 | -0.9 | 3.1 |
| *Can/Alc* | 2.3 | 1.0 | 2.2 | 0.031 | 0.2 | 4.3 |

**Model 2:** $K6 = X_0 + Substances + Age + Sex + Days\ from\ event$

Reference group: *No-Use*

| | |
|---|---|
| ***No. Observations*** | 434 |
| ***DoF Residuals*** | 424 |
| ***DoF Model*** | 9 |
| ***R-squared*** | 0.081 |
| ***Adj. R-squared*** | 0.061 |
| ***F-statistic*** | 4.134 |
| ***Prob (F-statistic)*** | 4.04e-05 |
| ***BIC*** | 2697 |

| | *Coefficient* | *SE* | *t* | *p* | *Lower CI* | *Upper CI* |
|---|---|---|---|---|---|---|
| *Intercept* | 10.8 | 1.3 | 8.3 | 0.0 | 8.2 | 13.3 |
| *Halluc* | 0.7 | 0.9 | 0.9 | 0.4 | -1.0 | 2.4 |
| *Halluc polyuse* | 0.8 | 0.8 | 1.1 | 0.3 | -0.7 | 2.4 |
| *MDMA* | -1.2 | 0.8 | -1.5 | 0.1 | -2.8 | 0.4 |
| *MDMA polyuse* | -0.3 | 0.8 | -0.4 | 0.7 | -1.9 | 1.3 |
| *Stim* | 1.0 | 1.0 | 1.0 | 0.3 | -1.0 | 3.0 |
| *Can/Alc* | 2.1 | 1.0 | 2.0 | 0.042 | 0.1 | 4.1 |
| *Age* | 0.06 | 0.04 | 1.7 | 0.096 | -0.011 | 0.1 |
| *Sex* | -2.0 | 0.5 | -3.9 | 0.0 | -3.0 | -1.0 |
| *Days from TE* | 0.01 | 0.01 | 1.2 | 0.2 | -0.01 | 0.03 |

**Model 1:** $\boldsymbol{PCL = X_0 + Substances}$

Reference group: *No-Use*

| | |
|---|---|
| ***No. Observations*** | 408 |
| ***DoF Residuals*** | 401 |
| ***DoF Model*** | 6 |
| ***R-squared*** | 0.04 |
| ***Adj. R-squared*** | 0.025 |
| ***F-statistic*** | 2.755 |
| ***Prob (F-statistic)*** | 0.0123 |
| ***BIC*** | 3408 |

| | *Coefficient* | *SE* | *t* | *p* | *Lower CI* | *Upper CI* |
|---|---|---|---|---|---|---|
| *Intercept* | 39.8 | 1.6 | 25.6 | 0.0 | 36.8 | 42.9 |
| *Halluc* | 4.7 | 2.7 | 1.7 | 0.1 | -0.6 | 9.9 |
| *Halluc polyuse* | 1.9 | 2.3 | 0.8 | 0.4 | -2.7 | 6.5 |

| | | | | | | |
|---|---|---|---|---|---|---|
| *MDMA* | -2.6 | 2.5 | -1.0 | 0.3 | -7.5 | 2.4 |
| *MDMA polyuse* | -1.0 | 2.4 | -0.4 | 0.7 | -5.7 | 3.8 |
| *Stim* | 3.5 | 3.1 | 1.1 | 0.3 | -2.5 | 9.5 |
| *Can/Alc* | 8.5 | 3.1 | 2.7 | 0.006 | 2.4 | 14.5 |

**Model 2:** $PCL = X_0 + Substances + Age + Sex + Days\ from\ event$

Reference group: *No-Use*

| | |
|---|---|
| ***No. Observations*** | 408 |
| ***DoF Residuals*** | 398 |
| ***DoF Model*** | 9 |
| ***R-squared*** | 0.08 |
| ***Adj. R-squared*** | 0.059 |
| ***F-statistic*** | 3.83 |
| ***Prob (F-statistic)*** | 0.0001 |
| ***BIC*** | 3408 |

| | *Coefficient* | *SE* | *t* | *p* | *Lower CI* | *Upper CI* |
|---|---|---|---|---|---|---|
| *Intercept* | 35.7 | 3.9 | 9.2 | 0.0 | 28.1 | 43.3 |
| *Halluc* | 5.7 | 2.6 | 2.1 | 0.033 | 0.5 | 10.9 |
| *Halluc polyuse* | 3.6 | 2.3 | 1.5 | 0.1 | -1.0 | 8.2 |
| *MDMA* | -1.3 | 2.5 | -0.5 | 0.6 | -6.2 | 3.6 |
| *MDMA polyuse* | 0.1 | 2.4 | 0.04 | 1.0 | -4.6 | 4.8 |
| *Stim* | 3.0 | 3.0 | 1.0 | 0.3 | -2.9 | 9.0 |
| *Can/Alc* | 8.1 | 3.0 | 2.7 | 0.008 | 2.1 | 14.1 |
| *Age* | 0.1 | 0.1 | 1.2 | 0.2 | -0.1 | 0.4 |
| *Sex* | -5.4 | 1.5 | -3.5 | 0.001 | -8.4 | -2.4 |
| *Days from TE* | 0.05 | 0.03 | 1.7 | 0.089 | -0.008 | 0.1 |

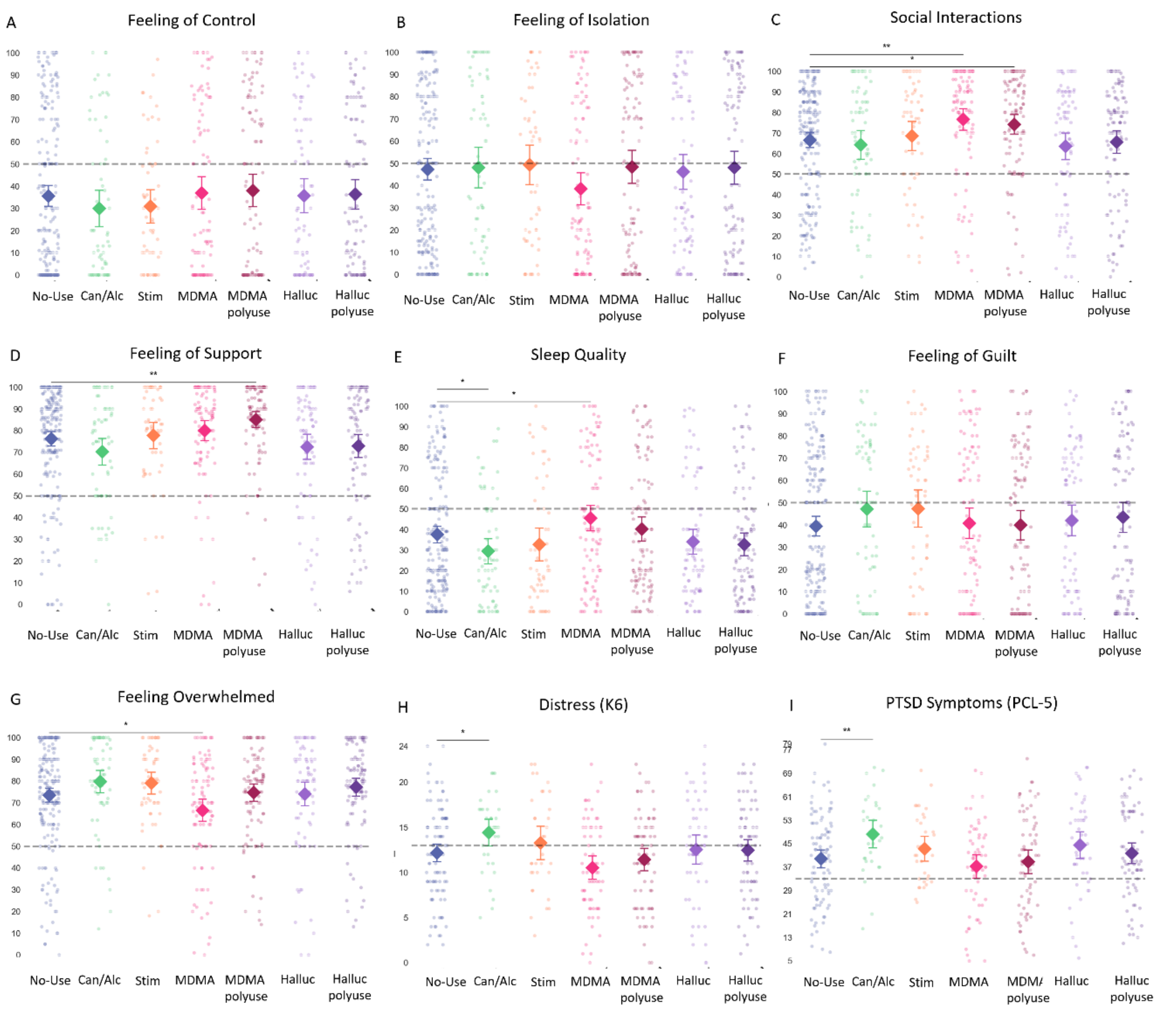


**Figure S2: Comprehensive Analysis Across All Substance Groups.** Panels A–I represent dot plots with overlaying mean values (diamonds) and 95% confidence intervals (error bars) for the respective outcome measures across substance use categories (*No-Use, Can/Alc, Stim, MDMA, MDMA Polyuse, Halluc, & Halluc Polyuse*). The following measures are included: (A) Feeling of Control; (B) Feeling of Isolation; (C) Social Interactions; (D) Feeling of Support (E) Sleep Quality; (F) Feeling of Guilt; (G) Feeling Overwhelmed; (H) Distress (K6); (I) PTSD Symptoms (PCL-5); Statistical significance is denoted by asterisks (*p < 0.05, **p < 0.01).

*Relative Risk*

The Relative Risk (RR) assessment was utilized to measure the influence of different substances on achieving favorable outcomes for the primary outcomes measures (Feeling Overwhelmed, K6, PCL-5). Supplementary Table S7 depict the RR for individuals in each group (numerator) compared to the *No-Use* group (denominator). An RR smaller than 1 suggests superior outcomes for the *No-Use* group.

**Table S7.** Relative Risk for favorable outcome against the *No-Use* group

| | *Overwhelmed* | *Mental distress (K6)* | *Posttraumatic symptoms (PCL-5)* |
|---|---|---|---|
| *Halluc* | RR=0.77<br>CI=[0.38,1.55]<br>P=0.47 | RR=0.89<br>CI=[0.63,1.25]<br>P=0.51 | RR=0.61<br>CI=[0.33,1.14]<br>P=0.12 |
| *Halluc polyuse* | RR=0.74<br>CI=[0.39,1.42]<br>P=0.36 | RR=0.85<br>CI=[0.62,1.16]<br>P=0.30 | RR=0.75<br>CI=[0.47,1.22]<br>P=0.25 |
| *MDMA* | RR=1.38<br>CI=[0.82,2.33]<br>P=0.23 | RR=1.22<br>CI=[0.94,1.59]<br>P=0.13 | RR=0.95<br>CI=[0.59,1.51]<br>P=0.81 |
| *MDMA polyuse* | RR=0.67<br>CI=[0.34,1.31]<br>P=0.24 | RR=1.00<br>CI=[0.75,1.34]<br>P=0.98 | RR=1.08<br>CI=[0.71,1.64]<br>P=0.72 |
| *Stim* | RR=0.37<br>CI=[0.12,1.18]<br>P=0.09 | RR=0.96<br>CI=[0.66,1.41]<br>P=0.85 | RR=0.71<br>CI=[0.37,1.39]<br>P=0.32 |
| *Can/Alc* | RR=0.74<br>CI=[0.34,1.61]<br>P=0.45 | RR=0.58<br>CI=[0.34,1.01]<br>P=0.05 | RR=0.28<br>CI=[0.09,0.84]<br>P=0.02 |

*Analyses of the Can/Alc group*

The *Can/Alc* group consists of survivors who consumed only Cannabis (n=16), only Alcohol (n=21) or both Cannabis and Alcohol (n=31). Due to the low number of survivors who used only Cannabis or only Alcohol, of which only a subset completed the full survey including the main outcome measures (Feeling overwhelmed: Cannabis=16, Alcohol=21; K6: Cannabis=9, Alcohol=14, PCL: Cannabis=9, Alcohol=13), and following reports from survivors indicating that Cannabis and Alcohol were considered to have a slighter phenomenological effect on one's state of consciousness compared to the other substance group, we combined these substances into one group in the main analyses. To validate this decision, we performed several analyses. First, we compared the Cannabis only vs. the Alcohol only groups and found no difference between them for the three outcome measures (All p-values>0.3). For each of the primary outcome measures (Feeling Overwhelmed, K6, PCL-5), a descriptive statistics table with the mean, standard deviation (SD), sample size (N), and confidence intervals (CI lower and upper) for each group is provided, followed by results from the linear regression models as follows:

- Model 1: The first model comparing the *Cannabis only* group to the *Alcohol only* group.
- Model 2: The second model adjusted for age, sex, and days since TE, comparing the *Cannabis only* group to the *Alcohol only* group.

*Feeling Overwhelmed*

| *Substances group* | *Mean* | *SD* | *N* | *ci lower* | *ci upper* |
|---|---|---|---|---|---|
| *Alcohol only* | 83.0 | 20.4 | 21 | 73.5 | 92.5 |
| *Cannabis only* | 81.3 | 17.5 | 16 | 71.7 | 90.9 |

**Model 1:** $\boldsymbol{Overwhelmed = X_0 + Substances}$

Reference group: *Alcohol Only*

| | |
|---|---|
| ***No. Observations*** | 37 |
| ***DoF Residuals*** | 35 |
| ***DoF Model*** | 1 |
| ***R-squared*** | 0.002 |
| ***Adj. R-squared*** | -0.027 |
| ***F-statistic*** | 0.06618 |

| | |
|---|---|
| ***Prob (F-statistic)*** | 0.798 |
| ***BIC*** | 331.0 |

| | *Coefficient* | *SE* | *t* | *p* | *Lower CI* | *Upper CI* |
|---|---|---|---|---|---|---|
| *Intercept* | 83.0 | 4.3 | 19.2 | 0.0 | 74.2 | 91.8 |
| *Cannabis only* | -1.7 | 6.6 | -0.3 | 0.8 | -15.0 | 11.6 |

**Model 2:**

$$\boldsymbol{Overwhelmed = X_0 + Substances + Age + Sex + Days\ from\ event}$$

Reference group: *Alcohol Only*

| | |
|---|---|
| ***No. Observations*** | 37 |
| ***DoF Residuals*** | 32 |
| ***DoF Model*** | 4 |
| ***R-squared*** | 0.028 |
| ***Adj. R-squared*** | -0.093 |
| ***F-statistic*** | 0.2320 |
| ***Prob (F-statistic)*** | 0.918 |
| ***BIC*** | 340.8 |

| | *Coefficient* | *SE* | *t* | *p* | *Lower CI* | *Upper CI* |
|---|---|---|---|---|---|---|
| *Intercept* | 79.5 | 20.9 | 3.8 | 0.0 | 37.0 | 122.0 |
| *Cannabis only* | -0.7 | 7.0 | -0.1 | 0.9 | -14.9 | 13.5 |
| *Age* | 0.2 | 0.6 | 0.3 | 0.8 | -1.0 | 1.4 |
| *Sex* | 4.4 | 7.1 | 0.6 | 0.5 | -9.9 | 18.8 |
| *Days from TE* | -0.1 | 0.1 | -0.7 | 0.5 | -0.3 | 0.2 |

*Mental Distress (K6 score)*

| *Substances group* | *Mean* | *SD* | *N* | *ci lower* | *ci upper* |
|---|---|---|---|---|---|
| *Alcohol only* | 12.8 | 3.9 | 14 | 10.5 | 15.1 |
| *Cannabis only* | 14.0 | 3.6 | 9 | 11.0 | 17.0 |

**Model 1:** $\boldsymbol{K6 = X_0 + Substances}$

Reference group: *Alcohol Only*

| | |
|---|---|
| ***No. Observations*** | 23 |
| ***DoF Residuals*** | 21 |
| ***DoF Model*** | 1 |
| ***R-squared*** | 0.024 |
| ***Adj. R-squared*** | -0.022 |
| ***F-statistic*** | 0.5166 |
| ***Prob (F-statistic)*** | 0.480 |
| ***BIC*** | 132.7 |

| | *Coefficient* | *SE* | *t* | *p* | *Lower CI* | *Upper CI* |
|---|---|---|---|---|---|---|
| *Intercept* | 12.8 | 1.1 | 12.1 | 0.0 | 10.6 | 15.0 |
| *Cannabis only* | 1.2 | 1.7 | 0.7 | 0.5 | -2.3 | 4.7 |

**Model 2:** $\boldsymbol{K6 = X_0 + Substances + Age + Sex + Days\ from\ event}$

Reference group: *Alcohol Only*

| | |
|---|---|
| ***No. Observations*** | 23 |
| ***DoF Residuals*** | 18 |
| ***DoF Model*** | 4 |
| ***R-squared*** | 0.065 |
| ***Adj. R-squared*** | -0.142 |
| ***F-statistic*** | 0.3142 |
| ***Prob (F-statistic)*** | 0.865 |
| ***BIC*** | 141.1 |

| | *Coefficient* | *SE* | *t* | *p* | *Lower CI* | *Upper CI* |
|---|---|---|---|---|---|---|
| *Intercept* | 14.7 | 4.8 | 3.0 | 0.007 | 4.6 | 24.9 |
| *Cannabis only* | 1.4 | 1.8 | 0.8 | 0.5 | -2.4 | 5.2 |
| *Age* | -0.1 | 0.1 | -0.7 | 0.5 | -0.4 | 0.2 |
| *Sex* | -0.6 | 1.8 | -0.3 | 0.7 | -4.4 | 3.2 |
| *Days from TE* | 0.02 | 0.03 | 0.6 | 0.6 | -0.1 | 0.1 |

*PTSD symptoms severity (PCL-5 scores)*

| *Substances group* | *Mean* | *SD* | *N* | *ci lower* | *ci upper* |
|---|---|---|---|---|---|
| *Alcohol only* | 42.5 | 11.5 | 13 | 35.3 | 49.7 |
| *Cannabis only* | 48.0 | 12.1 | 9 | 38.2 | 57.8 |

**Model 1:** $PCL = X_0 + Substances$

Reference group: *Alcohol Only*

| | |
|---|---|
| ***No. Observations*** | 22 |
| ***DoF Residuals*** | 20 |
| ***DoF Model*** | 1 |
| ***R-squared*** | 0.051 |
| ***Adj. R-squared*** | 0.004 |
| ***F-statistic*** | 1.081 |
| ***Prob (F-statistic)*** | 0.311 |
| ***BIC*** | 176.9 |

| | *Coefficient* | *SE* | *t* | *p* | *Lower CI* | *Upper CI* |
|---|---|---|---|---|---|---|
| *Intercept* | 42.5 | 3.4 | 12.5 | 0.0 | 35.4 | 49.6 |
| *Cannabis only* | 5.5 | 5.3 | 1.0 | 0.3 | -5.6 | 16.6 |

**Model 2:** $PCL = X_0 + Substances + Age + Sex + Days\ from\ event$

Reference group: *Alcohol Only*

| | |
|---|---|
| ***No. Observations*** | 22 |
| ***DoF Residuals*** | 17 |
| ***DoF Model*** | 4 |
| ***R-squared*** | 0.207 |
| ***Adj. R-squared*** | 0.021 |
| ***F-statistic*** | 1.110 |
| ***Prob (F-statistic)*** | 0.384 |
| ***BIC*** | 182.2 |

| | *Coefficient* | *SE* | *t* | *p* | *Lower CI* | *Upper CI* |
|---|---|---|---|---|---|---|
| *Intercept* | 27.5 | 14.6 | 1.9 | 0.1 | -3.3 | 58.3 |
| *Cannabis only* | 7.4 | 5.4 | 1.4 | 0.2 | -4.1 | 18.9 |

| | | | | | | |
|---|---|---|---|---|---|---|
| *Age* | 0.2 | 0.4 | 0.4 | 0.7 | -0.7 | 1.1 |
| *Sex* | 3.6 | 5.6 | 0.6 | 0.5 | -8.2 | 15.4 |
| *Days from TE* | 0.1 | 0.1 | 1.3 | 0.2 | -0.1 | 0.4 |

Furthermore, we compared the effects of alcohol vs. cannabis in conjunction with other substances (Cannabis polyuse n=68, Alcohol polyuse n=91). The results revealed, once again, no differences between these groups for the main outcome measures All p-values>0.6). For each of the primary outcome measures (Feeling Overwhelmed, K6, PCL-5), a descriptive statistics table with the mean, standard deviation (SD), sample size (N), and confidence intervals (CI lower and upper) for each group is provided, followed by results from the linear regression models as follows:

- Model 1: The first model comparing the *Cannabis polyuse* group to the *Alcohol polyuse* group.
- Model 2: The second model adjusted for age, sex, and days since TE,comparing the *Cannabis polyuse* group to the *Alcohol polyuse* group.

<u>*Feeling Overwhelmed*</u>

| *Substances group* | *Mean* | *SD* | *N* | *ci lower* | *ci upper* |
|---|---|---|---|---|---|
| *Alcohol polyuse* | 78.6 | 21.0 | 91 | 74.2 | 83.0 |
| *Cannabis polyuse* | 80.2 | 16.5 | 68 | 76.2 | 84.3 |

**Model 1:** $\boldsymbol{Overwhelmed = X_0 + Substances}$

Reference group: *Alcohol polyuse*

| | |
|---|---|
| ***No. Observations*** | 159 |
| ***DoF Residuals*** | 157 |
| ***DoF Model*** | 1 |
| ***R-squared*** | 0.002 |
| ***Adj. R-squared*** | -0.005 |
| ***F-statistic*** | 0.2714 |
| ***Prob (F-statistic)*** | 0.603 |
| ***BIC*** | 1402 |

| | *Coefficient* | *SE* | *t* | *p* | *Lower CI* | *Upper CI* |
|---|---|---|---|---|---|---|

| | | | | | | |
|---|---|---|---|---|---|---|
| *Intercept* | 78.6 | 2.0 | 38.7 | 0.0 | 74.6 | 82.6 |
| *Cannabis polyuse* | 1.6 | 3.1 | 0.5 | 0.6 | -4.5 | 7.7 |

**Model 2:**

$$\boldsymbol{Overwhelmed = X_0 + Substances + Age + Sex + Days\ from\ event}$$

Reference group: *Alcohol polyuse*

| | |
|---|---|
| ***No. Observations*** | 159 |
| ***DoF Residuals*** | 154 |
| ***DoF Model*** | 4 |
| ***R-squared*** | 0.088 |
| ***Adj. R-squared*** | 0.065 |
| ***F-statistic*** | 3.737 |
| ***Prob (F-statistic)*** | 0.00623 |
| ***BIC*** | 1402 |

| | *Coefficient* | *SE* | *t* | *p* | *Lower CI* | *Upper CI* |
|---|---|---|---|---|---|---|
| *Intercept* | 86.1 | 8.1 | 10.7 | 0.0 | 70.2 | 102.0 |
| *Cannabis polyuse* | 1.3 | 3.0 | 0.4 | 0.7 | -4.6 | 7.2 |
| *Age* | 0.2 | 0.2 | 0.7 | 0.5 | -0.3 | 0.6 |
| *Sex* | -9.5 | 3.2 | -3.0 | 0.003 | -15.7 | -3.2 |
| *Days from TE* | -0.1 | 0.1 | -1.8 | 0.077 | -0.2 | 0.01 |

*Mental Distress (K6 score)*

| *Substances group* | *Mean* | *SD* | *N* | *ci lower* | *ci upper* |
|---|---|---|---|---|---|
| *Alcohol polyuse* | 12.8 | 4.9 | 56 | 11.5 | 14.2 |
| *Cannabis polyuse* | 13.0 | 4.8 | 46 | 11.6 | 14.5 |

**Model 1:** $\boldsymbol{K6 = X_0 + Substances}$

Reference group: *Alcohol polyuse*

| | |
|---|---|
| ***No. Observations*** | 102 |
| ***DoF Residuals*** | 100 |
| ***DoF Model*** | 1 |
| ***R-squared*** | 0.001 |
| ***Adj. R-squared*** | -0.009 |
| ***F-statistic*** | 0.05111 |

| | |
|---|---|
| ***Prob (F-statistic)*** | 0.822 |
| ***BIC*** | 622.4 |

| | *Coefficient* | *SE* | *t* | *p* | *Lower CI* | *Upper CI* |
|---|---|---|---|---|---|---|
| *Intercept* | 12.8 | 0.7 | 19.4 | 0.0 | 11.5 | 14.1 |
| *Cannabis polyuse* | 0.2 | 1.0 | 0.2 | 0.8 | -1.7 | 2.2 |

**Model 2:**

$$K6 = X_0 + Substances + Age + Sex + Days\ from\ event$$

Reference group: *Alcohol polyuse*

| | |
|---|---|
| ***No. Observations*** | 102 |
| ***DoF Residuals*** | 97 |
| ***DoF Model*** | 4 |
| ***R-squared*** | 0.074 |
| ***Adj. R-squared*** | 0.035 |
| ***F-statistic*** | 1.929 |
| ***Prob (F-statistic)*** | 0.112 |
| ***BIC*** | 628.5 |

| | *Coefficient* | *SE* | *t* | *p* | *Lower CI* | *Upper CI* |
|---|---|---|---|---|---|---|
| *Intercept* | 15.4 | 2.5 | 6.1 | 0.0 | 10.4 | 20.4 |
| *Cannabis polyuse* | 0.2 | 1.0 | 0.2 | 0.8 | -1.7 | 2.1 |
| *Age* | -0.1 | 0.1 | -0.8 | 0.4 | -0.2 | 0.1 |
| *Sex* | -2.7 | 1.0 | -2.7 | 0.009 | -4.7 | -0.7 |
| *Days from TE* | 0.01 | 0.02 | 0.7 | 0.5 | -0.03 | 0.1 |

*PTSD symptoms severity (PCL-5 scores)*

| *Substances group* | *Mean* | *SD* | *N* | *ci lower* | *ci upper* |
|---|---|---|---|---|---|
| *Alcohol polyuse* | 43.3 | 13.1 | 53 | 39.7 | 46.9 |
| *Cannabis polyuse* | 42.7 | 15.6 | 43 | 37.9 | 47.6 |

**Model 1:** $PCL = X_0 + Substances$

Reference group: *Alcohol polyuse*

| | |
|---|---|
| ***No. Observations*** | 96 |
| ***DoF Residuals*** | 94 |
| ***DoF Model*** | 1 |
| ***R-squared*** | 0.000 |
| ***Adj. R-squared*** | -0.010 |
| ***F-statistic*** | 0.03871 |
| ***Prob (F-statistic)*** | 0.844 |
| ***BIC*** | 791.5 |

| | *Coefficient* | *SE* | *t* | *p* | *Lower CI* | *Upper CI* |
|---|---|---|---|---|---|---|
| *Intercept* | 43.3 | 2.0 | 21.9 | 0.0 | 39.4 | 47.2 |
| *Cannabis polyuse* | -0.6 | 3.0 | -0.2 | 0.8 | -6.4 | 5.3 |

**Model 2:**

$$PCL = X_0 + Substances + Age + Sex + Days\ from\ event$$

Reference group: *Alcohol polyuse*

| | |
|---|---|
| ***No. Observations*** | 96 |
| ***DoF Residuals*** | 91 |
| ***DoF Model*** | 4 |
| ***R-squared*** | 0.040 |
| ***Adj. R-squared*** | -0.003 |
| ***F-statistic*** | 0.9387 |
| ***Prob (F-statistic)*** | 0.445 |
| ***BIC*** | 801.3 |

| | *Coefficient* | *SE* | *t* | *p* | *Lower CI* | *Upper CI* |
|---|---|---|---|---|---|---|
| *Intercept* | 43.6 | 7.6 | 5.8 | 0.0 | 28.6 | 58.7 |
| *Cannabis polyuse* | 0.3 | 3.0 | 0.1 | 0.9 | -5.7 | 6.2 |
| *Age* | -0.1 | 0.2 | -0.7 | 0.5 | -0.6 | 0.3 |
| *Sex* | -3.5 | 3.1 | -1.1 | 0.3 | -9.6 | 2.7 |
| *Days from TE* | 0.1 | 0.1 | 1.6 | 0.1 | -0.02 | 0.2 |

Since no differences were found between individuals who consumed Cannabis and those who consumed Alcohol in the main outcome measures, we decided to collapse them into a single group to enhance statistical power.

*Additional variables of interest*

We collected additional variables to control for possible confounds between groups. Variables include therapy since the TE, substance experience prior to the TE, chronic medication use, neurological or psychiatric diagnoses, therapy before TE, and experience of previous trauma. For each variable, colored bars in Fig. S3 represents the proportion of individuals within each substance group that answered 'yes' to the question relative to the number of people that responded to the question, to account for differences in response rates between groups. Specifications regarding the response rates for each question per group are presented in Table S8.

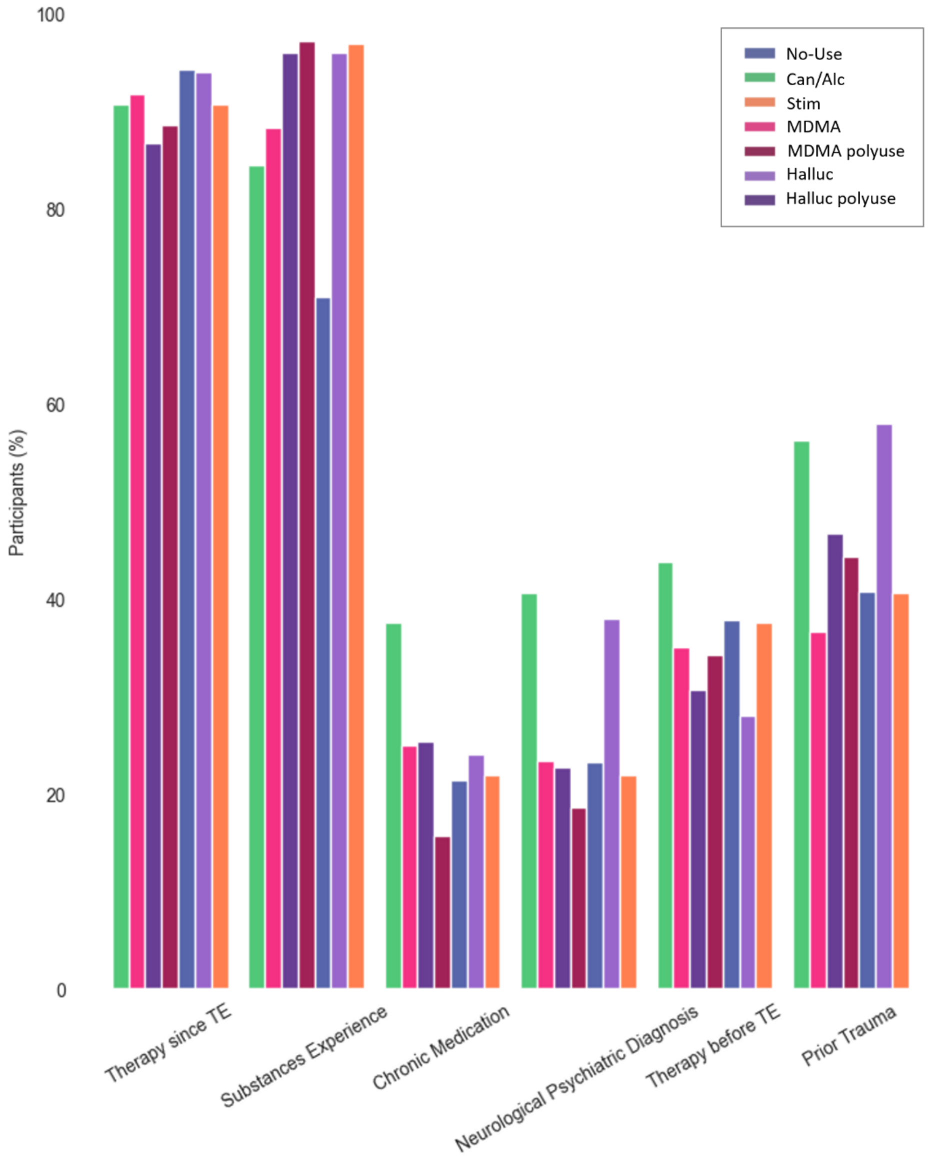


**Figure S3: Additional variables of interest.** Bar graph illustrating the percentages of participants reporting additional variables of interest, segmented by substance groups per legend. Variables include therapy since the TE, substance experience prior to TE, chronic medication use, neurological or psychiatric diagnoses, therapy before TE, and experience of previous trauma. Each bar represents the proportion of individuals within each substance group that answered 'yes' to the question relative to the number of people that responded to the question.

**Table S8**. Response distribution and demographics between participants in each of the substance groups (*Halluc*, *Halluc polyuse*, *MDMA*, *MDMA polyuse*, *Stim*, *Can/Alc*, *No-Use*). The table is organized to show both the total number of responses for each variable and the respective proportions within each group in parentheses, as not all participants answered every question. Most questions were optional. For example, for the question "Do you take chronic medications?", 104 participants from the *No-Use* group responded to the question, representing 48% of the entire group.

| | *Therapy since TE* | *Substance experience* | *Chronic medications* | *Neurological/ Psychiatric Diagnosis* | *Therapy before TE* | *Previous Trauma* |
|---|---|---|---|---|---|---|
| *Halluc* | 84 (100%) | 82 (98%) | 51 (61%) | 51 (61%) | 51 (61%) | 51 (61%) |
| *Halluc polyuse* | 107 (100%) | 104 (97%) | 77 (72%) | 77 (72%) | 77 (72%) | 77 (72%) |
| *MDMA* | 99 (100%) | 94 (95%) | 62 (63%) | 62 (63%) | 62 (63%) | 62 (63%) |
| *MDMA polyuse* | 108 (100%) | 103 (95%) | 71 (66%) | 70 (65%) | 71 (66%) | 71 (66%) |
| *Stim* | 58 (100%) | 56 (97%) | 34 (59%) | 34 (59%) | 34 (59%) | 34 (59%) |
| *Can/Alc* | 67 (99%) | 59 (87%) | 32 (47%) | 32 (47%) | 32 (47%) | 32 (47%) |
| *No-Use* | 216 (100%) | 199 (92%) | 104 (48%) | 105 (49%) | 104 (48%) | 104 (48%) |

*Correlation Analysis across substance groups*

As expected, the clinical measures of K6 and PCL-5 demonstrated a significant positive correlation (r=0.81; Fig. S4 panel A). These clinical measures were also positively correlated with the subjective feeling of being currently overwhelmed (K6 with overwhelmed: r=0.44; PCL with overwhelmed: r=0.49). Moreover, as per expectations, the social constructs were highly correlated with each other (Social support with Social interaction: r=0.48), affirming the intertwined nature of these social constructs. Interestingly, both social support and social interactions were inversely related to the primary clinical measures (Overwhelmed with social support: r=-0.20; Overwhelmed with social interactions: r=-0.23; K6 with social support: r=-0.37; PCL with social support: r=-0.34; K6 with social interactions: r=-0.42; PCL with social interactions: r=-0.39), suggesting that higher levels of perceived social support and frequent social interactions may be associated with better clinical outcomes. Sleep quality was also inversely correlated with the aforementioned clinical measures (sleep with Overwhelmed: r=-0.45; sleep with K6: r=-0.38; sleep with PCL: r=-0.51), suggesting that better sleep is associated with lower levels of psychological distress, PTSD symptoms severity and feelings of being overwhelmed. Moreover, feelings of guilt were positively correlated with K6 scores (r=0.47), PCL scores (r=0.48) and feeling overwhelmed (r=0.25).

*Correlation analysis by substance group*

When evaluating the correlation matrices across each substance group (Fig. S4 panels B-H), a notable observation is the moderate to strong positive correlations between Guilt and K6 scores and between Guilt and PCL-5 scores across all groups, indicating that individuals who feel more guilt also tend to report higher levels of distress and more severe PTSD symptoms following a TE. This relationship is consistent with clinical observations where guilt is often a symptom and a complicating factor in the recovery from TEs, potentially exacerbating the distress and severity of PTSD symptoms. Additionally, for the *Can/Alc* group, feelings of control show moderate positive correlations with feeling overwhelmed, distress (K6) and PTSD symptoms (PCL-5), while in all other groups these correlations are negative and weaker. This means that the more control individuals in the *Can/Alc* group felt during the TE, the guiltier they felt following the TE, in addition to having more severe psychological symptoms in the peritraumatic period. The reasons behind this unique correlation in the *Can/Alc* group could be numerous and require further investigation. It might reflect specific characteristics or experiences of the *Can/Alc* group, differences in how individuals in this group perceive or respond to control, or other underlying factors not directly measured in the study.

### Correlation Analyses

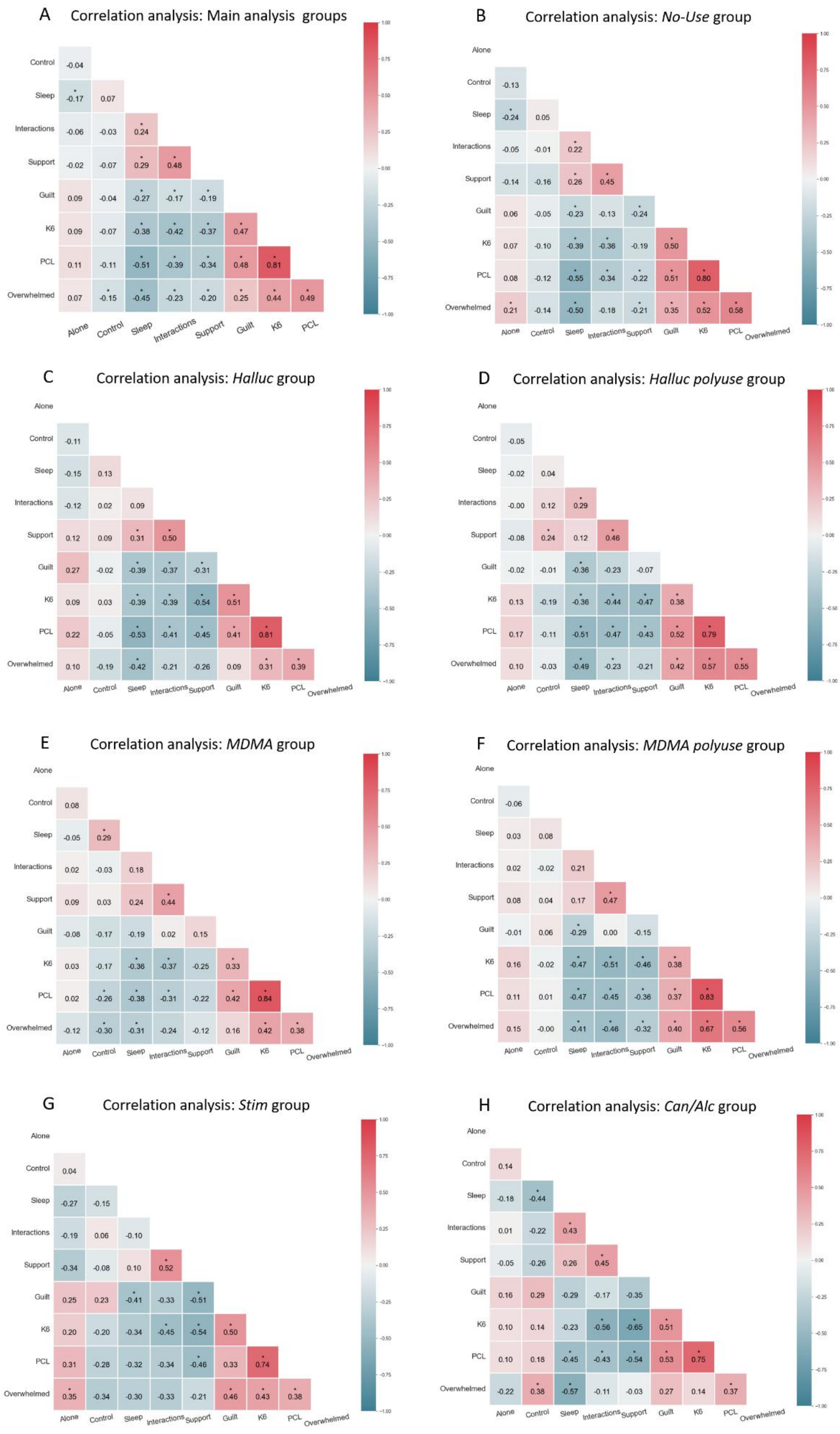

**Figure S4: Correlation analyses of outcome measures.** Correlation matrix heatmap showing the strength of associations between the different trauma related measures. Each matrix provides the Pearson correlation coefficients between outcome measures, including Isolation (*Alone*), Control, Sleep, Interactions, Support, Guilt, PCL-5 (posttraumatic symptoms), K6 (mental distress), and feelings of being Overwhelmed, to evaluate the interplay between different psychological domains. Correlation coefficients range from -1 to 1, with the scale illustrated on the right. Positive correlations are indicated in shades of red, negative correlations in blue, and neutral associations in white. Statistically significance correlations are denoted by asterisks (*p < 0.05). Panel A presents the correlation matrix across the four groups in the main analysis (*Halluc*, *MDMA*, *Can/Alc, No-Use*). Positive correlations are observed, particularly between clinical measures such as K6 and PCL scores (r = 0.81), as well as between these clinical measures and feeling overwhelmed and having feelings of guilt. K6 and PCL were found to be negatively correlated with social measures and sleep quality. Panels B-H present the correlation matrices for different substance groups: *No-Use* (B), *Halluc* (C), *Halluc polyuse* (D), *MDMA* (E), *MDMA polyuse* (F), *Stimulants* (G), *Can/Alc* (H).
